# TCF7L2 acts as a molecular switch in midbrain to control mammal vocalization through a transcriptional repression mechanism

**DOI:** 10.1101/2022.01.10.475593

**Authors:** Huihui Qi, Li Luo, Caijing Lu, Runze Chen, Xianyao Zhou, Xiaohui Zhang, Yichang Jia

**Author notes:** Corresponding author Please address correspondence to: Yichang Jia, Ph. D. School of Medicine, Medical Science Building, Room D204, Tsinghua University, Beijing, 100084, P. R. China. These authors contributed equally to this work.

## Abstract

Vocalization is an essential medium for sexual and social signaling in birds and mammals. Periaqueductal gray (PAG) a conserved midbrain structure is believed to be responsible for innate vocalizations, but its molecular regulation remains largely unknown. Here, through a mouse forward genetic screening we identified one of the key Wnt/ β-catenin effectors TCF7L2/TCF4 controls ultrasonic vocalization (USV) production and syllable complexity during maternal deprivation and sexual encounter. Expression of TCF7L2 in PAG excitatory neurons is necessary for the complex trait, while TCF7L2 loss reduces neuronal gene expressions and synaptic transmission in PAG. TCF7L2-mediated vocal β-catenin-binding domain but dependent of its DNA binding ability. Patient mutations associated with severe speech delay disrupt the transcriptional repression effect of TCF7L2, while mice carrying those mutations display severe USV impairments. Therefore, we conclude that TCF7L2 orchestrates gene expression in midbrain to control vocal production through a transcriptional repression mechanism.

## Introduction

Vocalization plays a fundamental role in intraspecific communication in songbirds, rodents, non-human primates and human (Nieder and Mooney, 2020). In songbirds and humans, some learning of vocal patterns, including songbird courtship call and human speech, are believed to rely heavily on imitation. In contrast, in rodent and non-human primate, vocalizations evoked by sexual cues or emotional states are considered to be largely innate. For example, mouse pups emit ultrasonic vocalizations (USVs) when separated from their mom, and adult male mice produces USVs in the presence of females. Such innate vocalization networks are located in the caudal brainstem and gated by the periaqueductal gray (PAG) region in the midbrain (Jurgens and Hage, 2007; Nieder and Mooney, 2020; Tschida et al., 2019). Importantly, the PAG to caudal brainstem circuit is highly conserved, appearing in both songbirds and humans, and is therefore believed to be required for vocal learning in these species. Recently, silencing of PAG-residing neurons that are transiently active during courtship vocalization revealed that these are both necessary and sufficient for USV production (Tschida et al., 2019). These finding suggest that deciphering the neural and molecular basis of the PAG to caudal brainstem circuit can help us understand both innate and learned vocalization mechanisms and their potential implications for human language disorders.

Human speech disorders have been linked with dysfunction of relatively few transcriptional factors (TFs) and their downstream genes (Hamdan et al., 2010; Horn et al., 2010; Konopka and Roberts, 2016; Lai et al., 2001; Newbury and Monaco, 2010). Point mutations of *FOXP2*, the most well-known vocalization gene, causes familiar language disorders and affected individuals exhibit deficits in production of words and complexity of syntax (Lai et al., 2001). Interestingly, misregulation of the *FOXP2* gene also impair mouse USV production (Castellucci et al., 2016; Chabout et al., 2016; Chen et al., 2016; Frohlich et al., 2017), underscoring common neuronal and molecular basis underlying human and mouse vocalization. To ensure it, we also generated *Foxp2* KO allele (Δ5) by Crispr/Cas9 approach and measured the pup USVs after maternal deprivation with a commercial USV detector (Figure S1). In agreement with the previous studies, *Foxp2*

KO dosage-dependently impaired the USV productions at the age points we examined. In contrast, the wildtype C57BL/6J littermates emitted reliable USVs especially at P5 and P7. In contrast to FOXP2, genetic alternations in Transcription Factor 7 Like 2 (*TCF7L2)*, the focus on the current study, is reportedly associated with human diseases, including metabolic disorders, cancers, and neurological disorders (Bem et al., 2019; Deciphering Developmental Disorders, 2015; Dias et al., 2021; Gonzalez et al., 2017; Grant et al., 2006; Iossifov et al., 2014; Stessman et al., 2017), but have not been reported in language disorders. Previous studies demonstrated that loss of TCF7L2 disrupts neuronal circuitry in a way that may explain the link between nicotine addiction and diabetes, and may play a role in the etiology of neuropsychiatric disorders (Duncan et al., 2019; Lipiec et al., 2020). TCF7L2 belongs to a family of T-cell factor/lymphoid enhancer (TCF/LEF) transcription factors (TCF7, LEF1, TCF7L1, and TCF7L2) that are key mediators of Wnt/ β-catenin signaling in numerous cellular processes ranging from early development to adult tissue homeostasis (Cadigan and Waterman, 2012; Clevers and Nusse, 2012; Hoppler and Kavanagh, 2007; Nusse and Clevers, 2017). TCF7L2 contains a β -catenin-mediated transcriptional activation and a high-mobility group (HMG) domain for DNA binding. In the absence of nuclear β catenin, a TCF7L2-containing transcriptional repression complex is assembled to inhibit downstream gene expression (Cadigan and Waterman, 2012; Daniels and Weis, 2005). Like other TCF/LEF family members, *TCF7L2* produces both full-length (flTCF7L2) and dominant-negative TCF7L2 (dnTCF7L2), the latter of which contains HMG box but not the CBD domain and therefore inhibits Wnt/ β-catenin signaling cascade in a dominant negative manner (Cadigan and Waterman, 2012; Hoppler and Kavanagh, 2007; Vacik and Lemke, 2011; Vacik et al., 2011).

Here, by using a N-ethyl-N-nitrosourea (ENU)-induced mutagenesis screening, we identified *Tcf7l2* acts as a gatekeeper for mouse USV production and syllable complexity. We show that *Tcf7l2* is highly expressed in midbrain VGLUT2 (vesicular glutamate transporter 2)-positive neurons and that its expression in these excitatory neurons is necessary and sufficiently for mouse USV production; midbrain region-specific knockout (KO) of *Tcf7l2* impairs mouse USV production and *Tcf7l2* KO in VGLUT2-positive neurons decreases synaptic transmission in PAG. Mice carrying patient mutations associated with developmental disorders display significant impairments of USV production. In addition, like the ENU-induced Y337H, patient non-synonymous mutations associated with severe speech delay relieve the transcriptional repression effect of TCF7L2. Therefore, we conclude that TCF7L2 functions as a molecular switch in the midbrain that controls vocalization production and syllable complexity.

## Results

### An ENU-induced mutagenesis screening for genes involved in mouse vocalization

To identify novel genes involved in mouse vocalization, we set up an ENU-induced mutagenesis screening (Figure 1A). To this end, we crossed ENU-treated G0 males with untreated C57BL/6J females, and through the use of an USV detector, identified G1 pups with a few or no USVs (< 5 times in a 5-minute interval at P5). Out of 702 G1 pups, 610 (86.9%) emitted USVs over 5 times whereas 92 (13.1%) emitted USVs 5 times or less, of which 13 mice (1.85%) were mute (Figure 1B and 1C). To establish family pedigrees of the ENU-induced mutations, we then crossed the adult USV-impaired G1 mice with untreated C57BL/6J mice to assess phenotypic reoccurrence in G2 and G3 pups. Among several families carrying inheritable USV impairments, we consistently found mute pups in the family #30 (Figure 1D and Figure S1).

**Figure 1.**
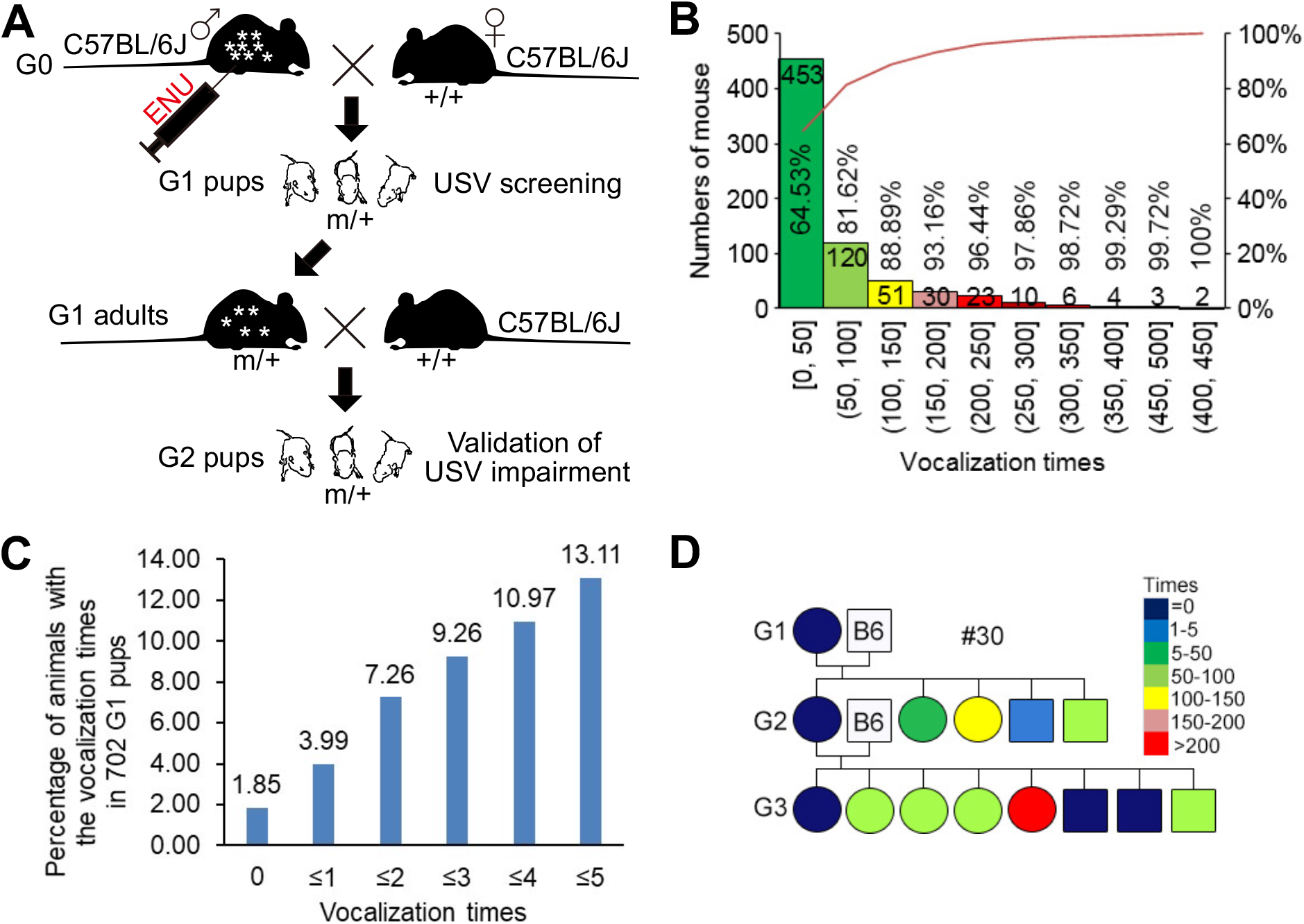
Identification of novel genes involved in mouse USV by an ENU- induced mutagenesis screening. (A) A G1 dominant screening was carried out for identification of novel genes involved in mouse USVs. G0 males were treated with ENU and then crossed to C57BL/6J wildtype (+/+) females. Spontaneous USVs of G1 pups induced by maternal deprivation were measured by a commercial USV detector (Med Associates Inc.). We screened for pups with USV impairments. The grownups of G1 pups with USV impairment were bred to C57BL/6J wildtype (+/+) mice. The reoccurrence of USV impairment in G2 and G3 pups was employed to establish the family pedigree. (B and C) The spontaneous USV number distribution of G1 pups (702) measured by the detector (Med Associates Inc.) in a 5-minute interval at P5. The G1 pups carried USV number ranged from 0-5 were illustrated (C). (D) An ENU family pedigree (#30 family) with an inheritable USV impairment at P5.

### ENU-induced Y337H mutation abolishes HMG box DNA binding ability

To identify the mutation causing the impaired USVs in the family #30, we conducted whole-exome sequencing of DNA from affected pups. Among the examined candidate mutations, we found that only a T to C non-synonymous mutation in the *Tcf7l2* gene (c.T1019C, p.Y337H, ENSMUST00000111656.7) is always co-segregated with the USV impairment (Figures 2A and S2). Notably, the T to C mutation changes the conserved 337 tyrosine, that has a hydrophobic side chain, into a positively charged histidine in the HMG box domain. The 337 tyrosine residue is located between two methionine residues that are proposed to be essential for HMG-mediated binding to the minor grove and bending the DNA double helix (Love et al., 1995) (Figure 2B). Specifically, the four aromatic residues Y12, W40, Y51, and Y52 in the HMG domain of LEF1 in murine form a hydrophobic core that stabilizes the HMG structure (Love et al., 1995). As illustrated in Figure 2B and 2C, the Y12, W40, Y51, and Y52 residues correspond to Y337, W365, Y376, and Y377 in TCF7L2. Furthermore, the corresponding residues are also present in two other TCF/LEF family members (Figure S3), underscoring the functional importance of these hydrophobic residues. Importantly, the ENU-induced Y337H mutation does not change TCF7L2 expression levels in mouse brain (Figure 2D).

**Figure 2.**
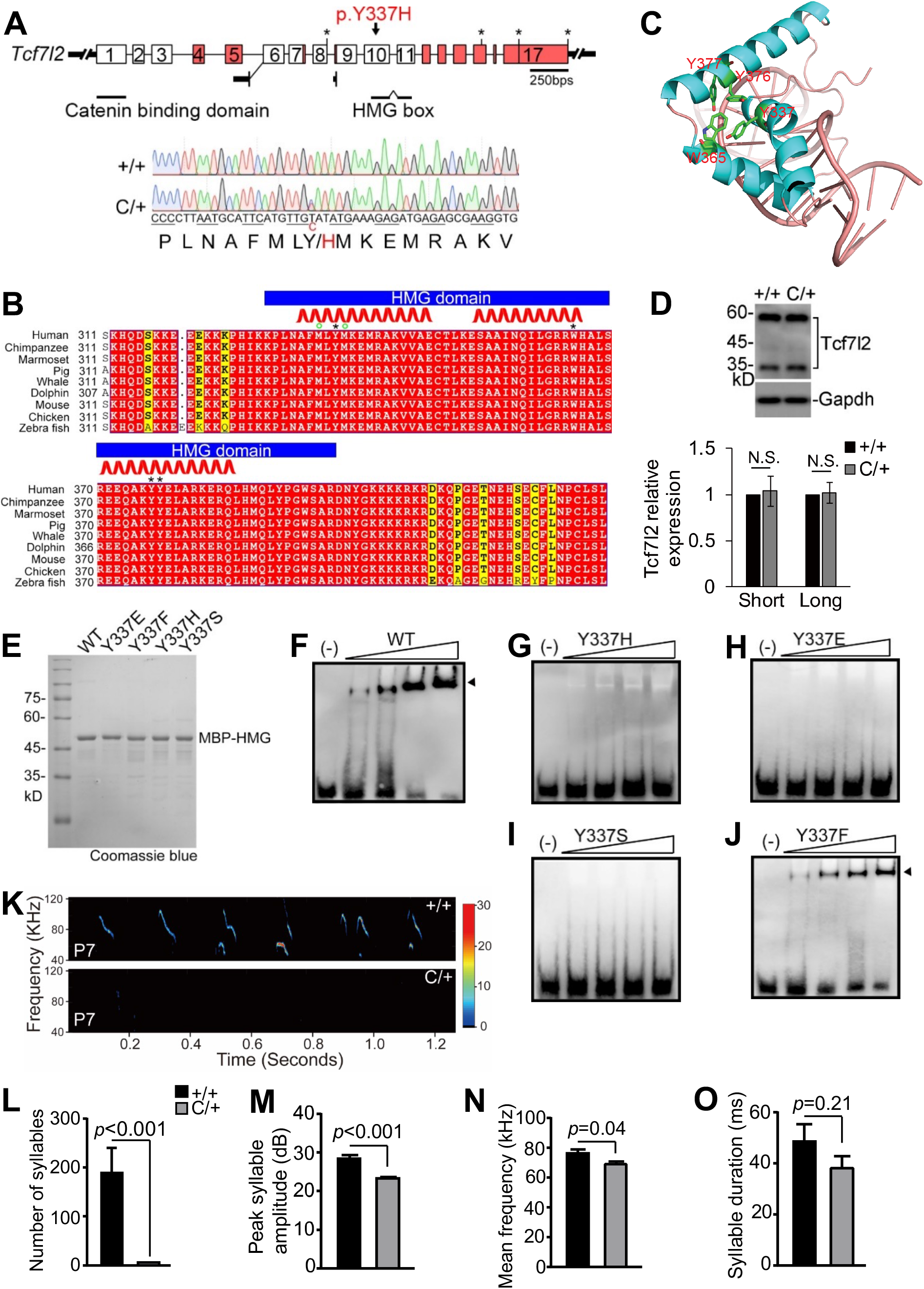
Y337H abolishes TCF7L2 DNA binding ability and impairs pup USVs. (A) The genome structure of mouse *Tcf7l2*. The alternatively spliced exons were marked in red. The ENU-induced nonsynonymous mutation identified in family #30 (c.T1019C, p.Y337H, ENSMUST00000111656.7) is located in *Tcf7l2* exon 10 that encodes part of HMG box. DNA chromatogram illustrates the mutation (C/+) (lower). (B) The sequence conservation of TCF7L2 HMG box and its neighboring residues. *, Y337, W365, Y376, and Y377; ^O^, M335 and M338; red ribbon, alpha helix segments. (C) The Y337 and neighboring residues were adapted into previously reported HMG box/DNA interface. The hydrophobic core formed by Y337, W365, Y376, and Y377 is highlighted. (D) The expression of TCF7L2 in wildtype (+/+) and mutant (C/+) thalamus. GAPDH, as a loading control. TCF7L2 appeared two major bands (Short and Long) in thalamus (upper) and statistic analysis of the relative expression level of these two bands (lower) in +/+ and C/+ mice. The anti-TCF7L2 antibody used here was generated by a synthetic peptide corresponding to sequences in HMG box. (E-J) HMG/DNA binding ability measured by EMSA. WT and the various mutant TCF7L2 HMG proteins fused with MBP-tag were purified by anti-MBP beads (E). (K) Representative spectrogram of USVs produced by +/+ and C/+ pups detected by MUPET. (L-O) The key features of USVs produced by +/+ and C/+ pups at P7. The value are presented as mean ± SEM. In D, n=4 (+/+ and C/+); in L-O, +/+, n=7, C/+, n=8; t-test, SPSS. N.S., no significant difference. See also Figure S2-S4.

W40 and Y51 mutations are known to impair LEF1 HMG DNA binding (Love et al., 1995); however, whether the Y12 mutation impacts on the DNA binding ability of the HMG domain of TCF/LEF family members is unknown. To answer this question, we mutated Y337 to: i) glutamic acid (E) to introduce a negative charge, ii) phenylalanine (F) to introduce a hydrophobic side chain), iii) histidine (H) to introduce a positive charge, and iv) serine (S) to introduce hydrophilic side chains (Figure 2E). In electrophoresis mobility shift assays (EMSA), we confirmed dose-dependent DNA binding of WT HMG (Figure 2F). However, we found that the HMG/DNA binding ability was abolished in Y337H as well as Y337E mutants (Figure 2G and 2H), suggesting that an impairment of HMG/DNA binding by Y337H is not due to a change in charge property. The HMG/DNA binding ability was also abolished in the Y337S mutant but largely retained by Y337F mutant HMG (Figure 2I-2J), suggesting that the hydrophobic property of Y337 is crucial for the HMG/DNA binding. Taken together, our data suggest that the ENU-induced Y337H mutation disrupts TCF7L2 binding to its target DNA, primarily by impairing formation of the HMG hydrophobic core domain.

### Y337H impairs pup vocalization and adult male vocal performances in a female-associated context

Animals heterozygous for the ENU-induced mutation (C/+) are viable and fertile with normal brain morphologies at P7 and 4 months of age (Figure S4A and S4B). As assessed by open field and Rotarod tests, C/+ mice also exhibit normal motor abilities relative to littermate wildtype controls (+/+) (Figure S4C-S4E). However, severe USV abnormalities are evident in mutant pups (Figure 2K-2O), including significantly fewer syllable numbers, lower peak amplitude, and reduced mean frequency, as detected by MUPET, an open-access USV analyzer (Van Segbroeck et al., 2017).

Distinct from pup vocalization, adult male and female mice emit USVs when they meet each other and the syllable complexity of such USVs is known to impact on the quality of male and female mouse social interactions (Chabout et al., 2015; Yang et al., 2013). Here, we reliably recorded male USVs when we put +/+ virgin males together with +/+ virgin females, irrespective of whether the females were anesthetized (AF) - and therefore unable to produce USVs upon meeting male mice - or live/awake (LF) (Figure 3A-3F). However, when we put virgin C/+ males with awake females, we detected significantly fewer syllable numbers, decreased peak amplitude, and shorter duration (Figure 3A-3F), indicative of vocal communication defects in a female-associated context. Relative to +/+ virgin males, the C/+ males also produced significantly fewer syllable numbers of shorter duration in the presence of anesthetized females. Taken together, we conclude that the Y337H mutation impairs generation of USVs in adult males upon meeting female mice.

**Figure 3.**
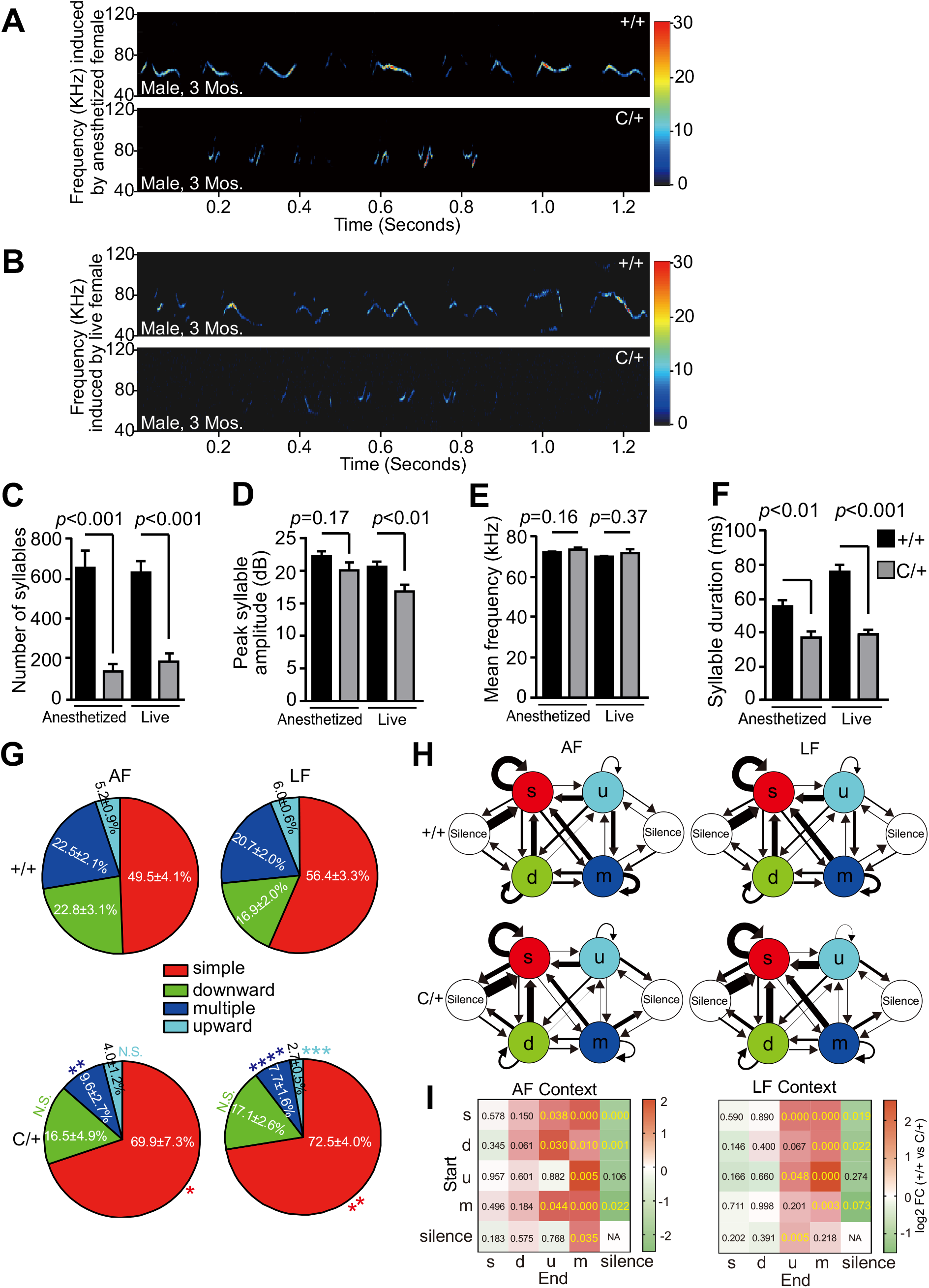
Y337H impairs adult male vocal performances in a female-associated context. (A and B) Representative spectrogram of USVs emitted by the virgin +/+ or C/+ male in the contexts of anesthetized (AF, A) or live/awake (LF, B) virgin female (+/+). Mouse age, 3 months. (C-F) The key features of USVs shown in (A) and (B). (G and H) Different repertoire compositions of syllables (G) and different conditional syllable transition probabilities (H) between +/+ and C/+ males with anesthetized (AF) or live/awake +/+ females (LF) shown in (A) and (B). In H, arrow direction and thickness represent the sequence and probability of syllable transition, respectively. Four syllable categories as previously described (PMID: 25883559): red, simple; green, downward; light blue, multiple; dark blue, upward. (I) Statistic analysis of conditional syllable transition probability between +/+ and C/+ virgin males. Number represents *p* value; color represents log2 value of the foldchange (FC). The value are presented as mean ± SEM. In C-F, G, and H, +/+, n=10, C/+, n=8; t-test, SPSS. N.S., no significant difference.

Next, we analyzed the syllables emitted by the C/+ or +/+ males in the presence of either awake or anesthetized females (Figure 3G). As previously described (Chabout et al., 2015), we categorized the syllables into 4 major types, including simple (s), multiple (m), downward (d), and upward (u). Relative to the +/+ virgin males, the C/+ males generated significantly more ‘s’ but fewer ‘m’ syllables in both settings. In addition, the C/+ males sang significantly fewer ‘u’ syllables in the presence of awake but not anesthetized females. Taken together, these data support that the Y337H mutation lowers the capability of generating syllable complexity in a female-associated context.

When we investigated the probability of mouse syllable transition, which reflects mouse ‘syntax’ to some degree (Chabout et al., 2015), we observed that fewer syllable transitions ended up with ‘m’ type in C/+ virgin males (Figure 3H). Moreover, more transitions ended up with ‘silence’ type in the mutant males, indicative of an early stop in the ‘syntax’ sequence. To quantify syllable transitions, we calculated transition probability from one type of syllable to all types with a fixed starting syllable. This approach avoids bias caused by extremely low numbers of syllable types, frequently seen in the C/+ animals (Chabout et al., 2015). In the presence of anesthetized females, we found that both syllable transition probabilities starting with any types and ending up with ‘m’ and starting with ‘s’, ‘d’, and ‘m’ types and ending up with ‘u’ were significantly higher in +/+ than in C/+ virgin males (Figure 3H and 3I). Similarly, in the presence of awake females, syllable transition probabilities from ‘s’, ‘d’, ‘u’, and ‘m’ to ‘m’ and from ‘s’, ‘u’, and ‘silence’ to ‘up’ category were significantly higher in +/+ than in C/+ virgin males (Figure 3H and 3I). In contrast, in both contexts, the transition probabilities from ‘s’, ‘d’, and ‘m’ to ‘silence’ were significantly lower in +/+ than in C/+ males. Based on these data, we conclude that the Y337H mutation impairs ability to complete specific syllable transitions, which breaks ‘syntax’ sequence continuity and causes premature stop.

### Haploinsufficiency of *Tcf7l2* impairs pup vocalization and reduces female preference

The Y337H mutation does not affect TCF7L2 expression (Figure 2D) but abolishes DNA binding ability (Figure 2G), suggesting that TCF7L2 carrying the Y337H mutation impairs mouse vocalization through a loss-of-function mechanism. Next, we generated a *Tcf7l2* KO mouse model by using Crispr/Cas9 technology to introduce a two-nucleotide deletion (Δ2) into *Tcf7l2* exon 10 (Figure 4A and 4B). Heterozygous *Tcf7l2* KO mice (Δ2/+) are viable and fertile with reduced expression of high and low molecular weight TCF7L2 in the midbrain (Figure 4C). Similar to the ENU-induced Y337H mutation (Figure 2K-2O), *Tcf7l2* haploinsufficiency severely impaired USV generation (including significantly less syllable numbers, peak amplitude, and mean frequency) relative to control animals (Figure 4D-4G).

**Figure 4.**
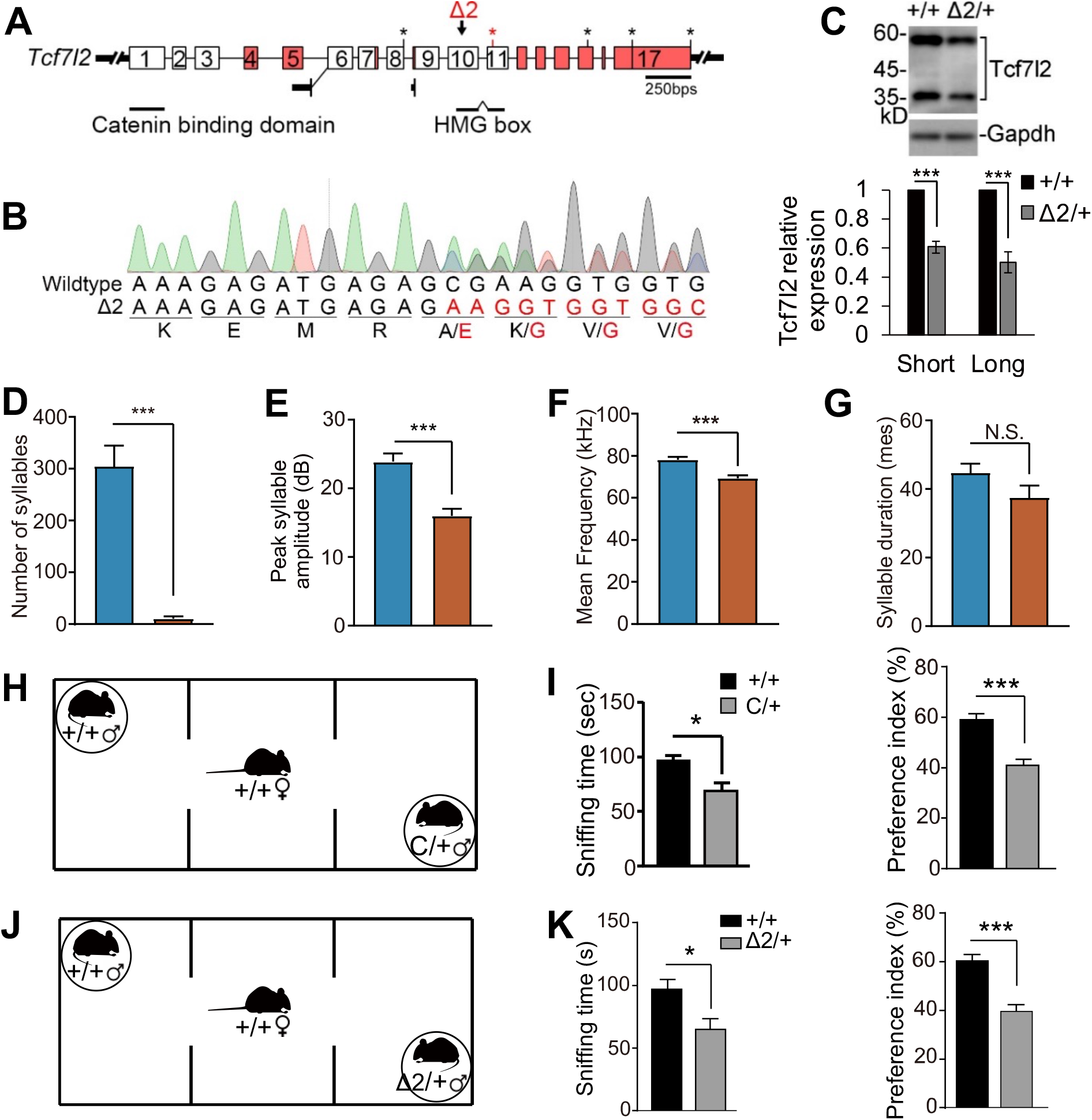
Haploinsufficiency of *Tcf7l2* impairs pup USVs and reduces sexual preference. (A and B) A 2-nucleotide deletion (Δ2) in *Tcf7l2* exon 10, which encodes part of TCF7L2 HMG domain, was introduced by Crispr/Cas9. The deletion leads to a premature termination codon (red asterisk) in *Tcf7l2.* DNA chromatogram (B) of the heterozygous KO mouse (Δ2/+). (C) The Δ2 deletion significantly reduces the expression of TCF7L2 in midbrain. GAPDH as loading control. Short, low molecular weight (∼35kD) TCF7L2; Long, high molecular weight (∼60kD) TCF7L2. (D-G) The key features of USVs produced by the wildtype (+/+) and KO mutant (Δ2/+) pups at P7. (H-K) Haploinsufficiency of *Tcf7l2* in male reduces female preference. Diagrams illustrates the female preference measurement by modification of three-chamber test (H and J). The data summary was shown in (I and K). The value are presented as mean ± SEM. In C, n=3; in D-G, +/+ (n=16), Δ2/+ (n=9); in I, +/+ (male, n=7), +/+ (female, n=7), C/+ (male, n=7); in K, +/+ (male, n=7), +/+ (female, n=7), Δ2/+ (male, n=7). N.S., no significant difference, * *p*<0.05, *** *p*<0.001, t-test, SPSS.

Because the Y337H mutation in the ENU model lowers male syllable complexity and breaks their ‘syntax’ sequence continuity (Figure 3), we asked whether this vocalization defect would cause wildtype female mice to prefer wildtype male mice. To answer this question, we modified a three-chamber test by restricting virgin male mice to two separate sides and allowing virgin female to freely access the two males and show their preference, as previously described (Tschida et al., 2019) (Figure 4H, 4J). In this setting, we found that the female mice spent significantly more sniffing time with the wildtype males relative to both C/+ and Δ2/+ male mice (Figure 4I,4K). In addition, based on the preference index, females also preferred spending time with wildtype over both C/+ and Δ2/+ male mice (Figure 4I,4K). Taken together, these studies show that male *Tcf7l2* haploinsufficiency reduces female preference.

### Expression of TCF7L2 in *Vglu2*-positive neurons is sufficient and necessary for mouse vocalization

Through immunostaining and *in situ* hybridization analysis, we demonstrate that TCF7L2 expression is concentrated in mouse thalamus and midbrain at P7 (Figure S5A,B) consistent with a previous report (Nagalski et al., 2013), and that *Tcf7l2*-expressing cells are *Vglut2*-positive excitatory neurons (Figure S5B and S5C). Quantitative-PCR and immunoblot analysis further confirmed that *Tcf7l2* mRNA and TCF7L2 protein expression is concentrated in thalamus and midbrain at P7 (Figure S5D and S5E). In all age points examined, we detected two molecular weight TCF7L2 (Short, ∼35kD; Long, ∼60kD), both of which were significantly reduced in *Tcf7l2* heterozygous KO (Δ2/+) midbrain (Figure 4C). Next, we employed a previously reported *Tcf7l2* flox allele, in which the loxP sites flank exon 11 that partly encodes for the HMG box (van Es et al., 2012), to determine which *Tcf7l2*-expressing cell type is responsible for mouse vocalization. Similarly to what we observed in the Δ2 mouse, midbrain expression of both short and long forms of TCF7L2 were reduced in *exon11 fl*/+;*Nestin-Cre*/+ mice (Figure S6A). Consequently, USVs in *exon11 fl*/+;*Nestin-Cre*/+ P7 pups displayed significantly fewer syllables, reduced peak amplitude, lower mean frequency, and shorter duration than controls (Figure 5A), indicating that *Tcf7l2* expression in neuronal progenitor is required for pup USVs.

**Figure 5.**
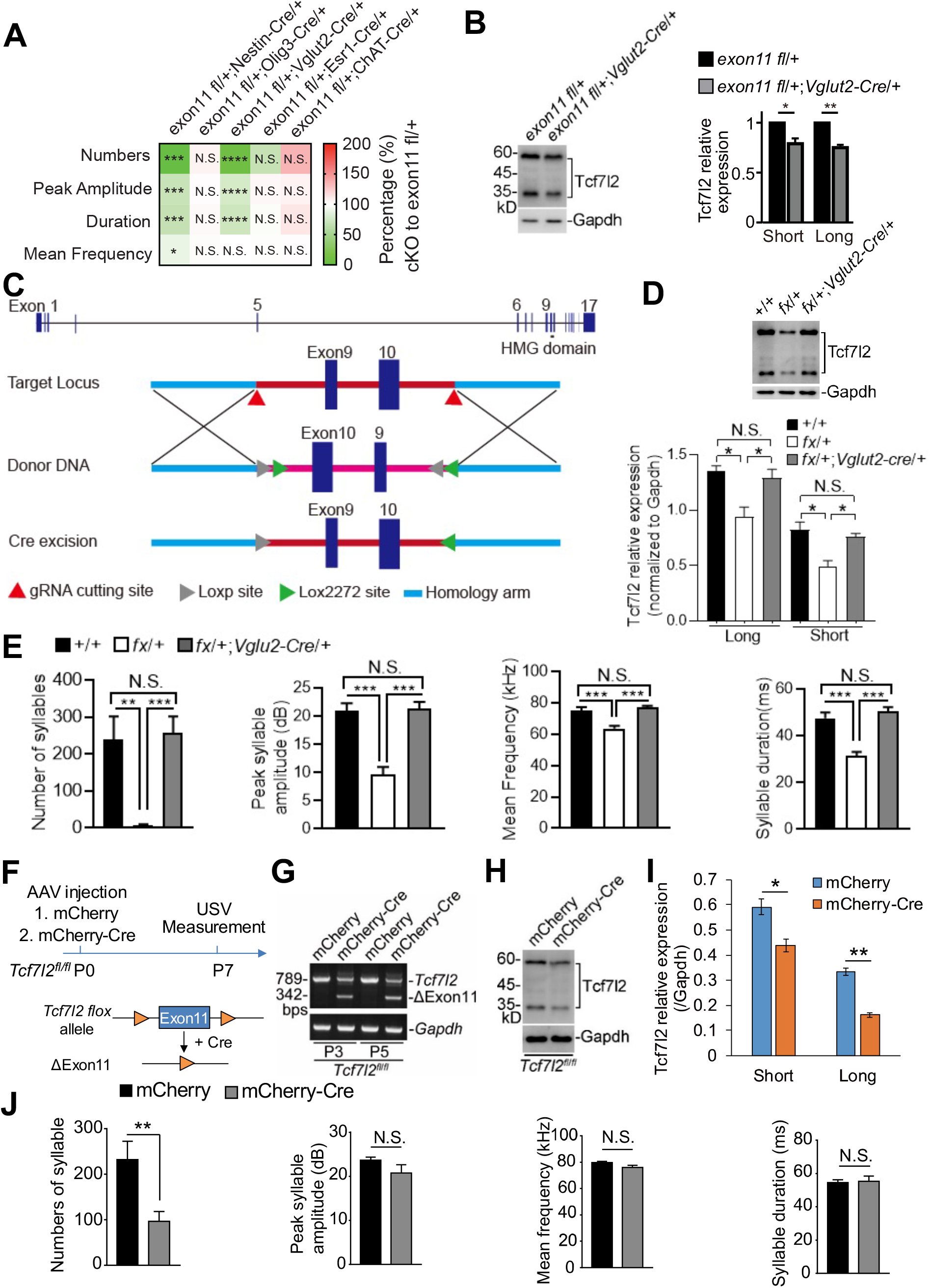
Expression of *Tcf7l2* in *Vglut2*-positive neurons in midbrain is necessary and sufficient for mouse USV production. (A) The key features of pup USVs produced by the indicated *Tcf7l2* cKO pups at P7. *Nestin-*, *Oligo3-*, *Vglut2*-, *Esr1*-, and *ChAT*-*Cre* were employed to conditionally remove *Tcf7l2*. (B) The expression level of TCF7L2 in the midbrains of *exon11 fl*/+;*Vglut2*-*Cre*/+ mice by western blot. The *exon11 fl*/+ mice served as controls; GAPDH, loading control. (C) A *Cre*-dependent FLEx switch allele (fx) containing an inverted *Tcf7l2* exon 9 and 10, which encode part of HMG domain, was flanked with inward-facing tandem Lox sites (LoxP:Lox2272). (D) Expression level of TCF7L2 in midbrain in the indicated genotypes at P7. In the absence of Cre recombinase, the expression level of TCF7L2 was significantly reduced in the *fx*/+ mice, which was restored by crossing to *Vglut2*-*Cre*. (E) Cre expression in *Vglut2*-positive neurons restores the pup USVs shown in the *fx*/+;*Vglut2*-*Cre*/+ mice at P7. (F) A schematic diagram for *Tcf7l2* KO in midbrain by AAV-mCherry-Cre injection. The AAV viral particles were injected into midbrain of *Tcf7l2 exon11 fl*/*fl* pups at P0 and USVs were measured at P7. The Cre expression was driven by a neuronal Syn1 promoter. AAV-mCherry served as a control. (G) The removal of exon 11 (ΔExon11) at DNA level was detected in midbrain at P3 and P5 after AAV-mCherry-Cre injection. *Gapdh* served as a control for genomic DNA PCR. (H and I) Expression of TCF7L2 in PAG of *Tcf7l2 exon11 fl*/*fl* mice at P7 after AAV- mCherry or AAV-mCherry-Cre injections. Data summary was shown in (I) (n=3). (J) Pup USVs were measured in *Tcf7l2 exon11 fl*/*fl* mice injected with AAV-mCherry (n=9) and AAV-mCherry-Cre (n=12). In A, B, D, E, I, and J, data are presented as mean ± SEM. N.S., no significant difference, * *p*<0.05, ** *p*<0.01, *** *p*<0.001, t-test or ANOVA, SPSS. In A, *Tcf7l2 exon11 fl*/+ mice were employed as controls (n=7-15); *Tcf7l2 exon11 fl*/+;*Nestin*- *Cre*/+ (n=12); *Tcf7l2 exon11 fl*/+;*Oligo3*-*Cre*/+ (n=8); *Tcf7l2 exon11 fl*/+;*Vglut2*-*Cre*/+ (n=13); *Tcf7l2 exon11 fl*/+; *Esr1*-*Cre*/+ (n=7); *Tcf7l2 exon11 fl*/+;*ChAT*-*Cre*/+ (n=9). In B and D, n=3. In E, +/+ (n=12); *Tcf7l2 fx*/+ (n=12); *Tcf7l2 fx*/+;*Vglut2*-*Cre*/+ (n=9). See also Figure S5-S7.

Because *Tcf7l2* mRNA and TCF7L2 protein are highly expressed in the thalamus, we next asked whether thalamic expression of *Tcf7l2* is responsible for mouse vocalization. To this end, we generated *exon11 fl*/+;*Olig3-Cre*/+ mice, in which the cKO of *Tcf7l2* is achieved by the thalamic *Olig3-Cre* driver (Vue et al., 2009). Although expression of both short and long TCF7L2 were reduced in the thalamus of *exon11 fl*/+;*Olig3-Cre*/+ mice (Figure S6B) and the thalamic *Olig3-Cre* expression were confirmed (Figure S6C), USVs were normal (Figure 5A), indicating that expression of *Tcf7l2* in the thalamus *per se* is not a requirement for pup USVs.

Given that *Tcf7l2*-expressing neurons are *Vglut2*-positive, we next generated *exon11 fl*/+;*Vglut2*-*Cre*/+ mice and validated that levels of both short and long TCF7L2 were reduced in midbrain in these mice (Figure 5B). Relative to P7 *exon11 fl*/+ control mice, we detected significantly fewer syllable numbers, lower peak amplitude, and shorter duration in *exon11 fl*/+;*Vglut2*-*Cre*/+ pups (Figure 5A), indicating that expression of *Tcf7l2* in *Vglut2*-positive neurons is necessary for pup vocalization. To examine whether *Tcf7l2* expression in *Vglut2*-positive neurons is also sufficient for mouse vocalization, we generated a *fx* allele for *Tcf7l2* (Figure 5C), which inverts exon 9 and 10 and places the inward-facing Lox2272:LoxP sites flanking the two inverted exons (Figure 5C). With this approach, we could switch on *Tcf7l2* expression in a Cre-dependent manner (Schnutgen et al., 2003). In the absence of *Cre*, we validated that the expression of both short and long TCF7L2 was significantly reduced in the midbrain of the *fx*/+ mice; however, the expression of TCF7L2 was restored in the midbrain of *fx*/+;*Vglut2*-*Cre*/+ mice (Figure 5D). Importantly, the pup USV abnormalities present in the *fx*/+ mice were absent in *fx*/+;*Vglut2*-*Cre*/+ mice (Figure 5E).

Recent work revealed that estrogen receptor 1 (ESR1) -expressing neurons in the preoptic area and the ventromedial hypothalamus participate in mouse USV (Chen et al., 2021; Gao et al., 2019; Karigo et al., 2021; Michael et al., 2020). In addition, motor neuron pools in the brain stem and spinal cord that control laryngeal and respiratory muscles also contribute to vocalization in mammals (Arriaga et al., 2012; Jurgens, 2002; Jurgens and Hage, 2007). Here, we found that pups derived from *exon11 fl*/+ mice crossed with *Esr1*-*Cre*/+ or *ChAT*-*Cre*/+ mice emitted normal syllable numbers, peak amplitude, mean frequency, and duration at P7 (Figure 5A). These experiments support that TCF7L2 expression in *Esr1*- or *ChAT*-positive neurons is not required for USV generation, but that expression of TCF7L2 in *Vglut2*-positive neurons is both necessary and sufficient for mouse vocalization.

### Midbrain expression of *Tcf7l2* is required for mouse vocalization

Although *Tcf7l2* expression is concentrated in mouse thalamus and midbrain (Figure S5), thalamic expression of *Tcf7l2* does not contribute to mouse USV production (Figure 5A), prompting us to hypothesize that *Tcf7l2* expression in PAG is responsible for mouse USV production. Indeed, we found that TCF7L2 is expressed in dorsal and lateral but not ventral PAG (Figure S7A) and that TCF7L2-positive cells were also positive for *Vglut2-Cre*-driven RPL22-HA expression (Figure S7B and S7C; dorsal 53.9%, lateral 75.2%), indicating that TCF7L2-positive cells in PAG are excitatory neurons.

Although TCF7L2 expression is present in the embryonic mouse brain from E12.5, and high in the midbrain between P0 and P7, upon maternal deprivation, pup USV production does not peak until P5 or P7 (Figure S1, S5). To examine whether midbrain *Tcf7l2* expression is responsible for USV production, we injected AAV-mCherry or AAV-mCherry-Cre into the midbrain of *exon11 fl*/*fl* pups at P0 and measured USV at P7 (Figure 5F). We found that the injection of AAV-mCherry-Cre but not AAV-mCherry removed exon 11 (Figure 5G) in the midbrain as early as P3, three days after the injection, and significantly reduced both short and long TCF7L2 in the midbrain at P7 (Figure 5H and 5I). In our USV production analysis, we only included pups with mCherry fluorescence at PAG, and found that, relative to AAV-mCherry injection, the AAV-mCherry-Cre injection significantly decreased syllable numbers and reduced both peak amplitude and mean frequency (Figure 5J). These data suggest that *Tcf7l2* expression in midbrain PAG *Vglut2*-positive neurons is responsible for mouse USV production.

### Full-length of TCF7L2 in *Vglu2*-positive neuron is required for mouse USV production

The TCF7L2 HMG domain is encoded by *Tcf7l2* exon 10 and 11. Because our anti-TCF7L2 antibody was generated from synthetic polypeptide corresponding to sequences in the HMG, and detects both high and low molecular weight bands (Figure S5E), we conclude that both the long and short form of TCF7L2 contains the HMG region. Indeed, both two-nucleotide deletion in exon 10 (Δ2/+) and conditional removal of exon 11 in *Vglut2*-positive neurons (*exon11 fl*/+;*Vglut2*-*Cre*/+) lead to reduction of both short and long TCF7L2 proteins (Figure 4C and 5B) and impaired USV (Figure 4 and Figure 5A). Next, we asked which form of TCF7L2 contributes to mouse USV production. To this end, we imported a previously reported *Tcf7l2* exon1 flox allele (*Tcf7l2 exon1 fl*), in which the CBD-encoding exon 1 is flanked by two loxP sites (Angus-Hill et al., 2011). We found that conditional removal of exon1 in *Vglut2*-positive neurons (*exon1 fl*/+;*Vglut2*-*Cre*/+) significantly impaired pup USVs compared to *exon1 fl*/+ controls (Figure 6A) and significantly reduced the long but not the short TCF7L2 form in the midbrain (Figure 6B and 6C), indicating that flTCF7L2 is required for mouse USV production.

**Figure 6.**
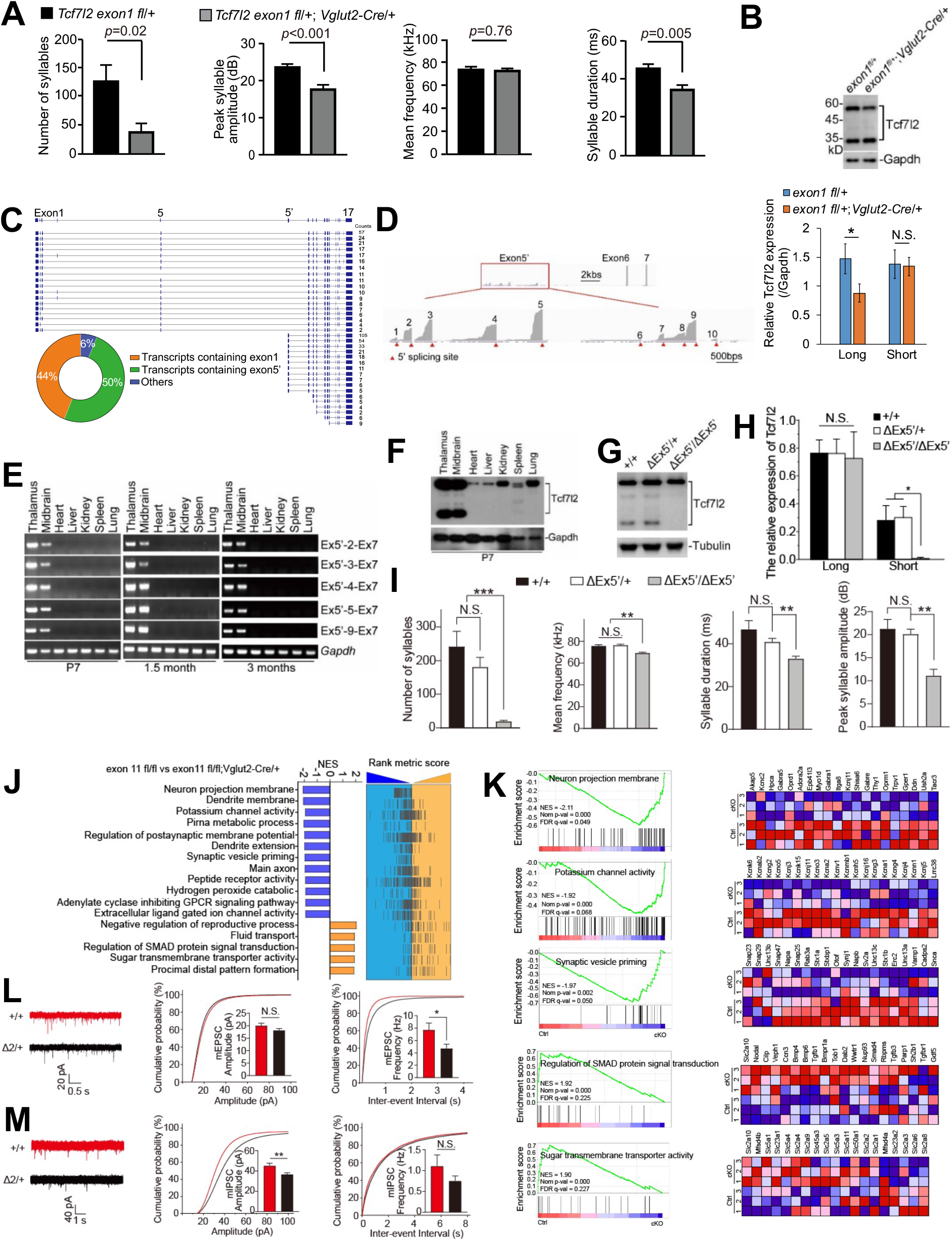
Both flTCF7L2 and dnTCF7L2 are required for mouse USVs and synaptic transmission in PAG is impaired by TCF7L2 loss. (A) USV measurement in *exon1 fl*/+;*Vglut2*-*Cre*/+ (n=15) at P7. The *exon1 fl*/+ mice (n=19) served as controls. (B) The expression level of TCF7L2 in the midbrain of *exon1 fl*/+;*Vglut2*-*Cre*/+ and *exon1 fl*/+ mice. GAPDH, loading control. Data summary shown in C (n=4) (C) Different isoforms of *Tcf7l2* in mouse brain was identified by nanopore sequencing and the isoform percentages were illustrated. (D) IGV visualization of 10 alternative exon 5’. Red triangles illustrate the 5’ splicing sites of these alternative exon 5’. (E) Transcripts containing exon 5’ were detected in neuronal but not non-neuronal tissues by RT-PCR at P7, 1.5-month, and 3-month-old mice. (F) DnTCF7L2 are expressed in neuronal but not non-neuronal tissues at P7. (G and H) Expression of dnTCF7L2 in the midbrains in the indicated genotypes at P7 (G). The data summary was shown in (H) (n=3). (I) USV measurement in the indicated genotypes at P7. (J) Summary of GSEA analysis in PAGs at P7 between *exon11 fl*/*fl* and *exon11 fl*/*fl*;*Vglut2*-*Cre*/+ mice. NES, normalized enrichment score. (K) Significantly enriched biological pathways and their individual genes identified by GSEA analysis. In heatmap, blue and red, downregulated and upregulated expression in the cKO PAGs, respectively. (L and M) Representative mEPSC and mIPSC trace recording from LPAG of Δ2/+ and +/+ mice at one-month of age and statistic analysis of the amplitudes and frequencies. In A, B, H, I, L, and M, the value are presented as mean ± SEM. N.S., no significant difference, * *p*<0.05, ** *p*<0.01, *** *p*<0.001, t-test or ANOVA, SPSS. In I, +/+ (n=10), ΔEx5’/+ (n=19), ΔEx5’/ΔEx5’ (n=9). In L and M, for mEPSC, +/+, n=21/3; Δ2/+, n=23/3; for mIPSC, +/+, n=22/4; Δ2/+, n=29/3. See also Figure S8 and S9.

### Brain-specific dnTCF7L2 is required for mouse vocalization

Our nanopore sequencing, a long-read sequencing technology (Venkatesan and Bashir, 2011), revealed that about 44% of *Tcf7l2* transcripts in mouse brain contain exon1 (Figure 6C). However, of all transcripts, about 50% contain exon 5’ and the following exons but not upstream exon 1-5, indicating that TCF7L2^short^ is dnTCF7L2 without the CBD (Cadigan and Waterman, 2012; Hoppler and Kavanagh, 2007; Vacik and Lemke, 2011; Vacik et al., 2011). When we took a closer look at exon 5’, we noticed that at least 10 alternative exon 5’ with 5’ splicing sites are expressed in mouse brain (Figure 6D). Among these 5’ exons, exon 5’-2, -3, -4, -5, and -9 were resolved by more nanopore reads. Furthermore, our RT-PCR analysis confirmed expression of these transcripts in midbrain and thalamus at P7, 1.5-, and 3-month of age (Figure 6E). However, the transcripts were absent in non-neuronal tissues, including heart, liver, kidney, spleen, and lung, suggesting that these transcripts are specific to neuronal tissue. Indeed, dnTCF7L2 proteins were detected in thalamus and midbrain but not in non-neuronal tissues (Figure 6F).

To validate that dnTCF7L2 is translated from exon 5’-containing transcripts, we removed exon 5’-2-9 by Crispr/Cas9-mediated genome editing (Figure S8A) and verified that expression of transcripts containing exon 5’-2, -3, -4, -5, and -9 was not detected in thalamus and midbrain in exon 5’ KO mice (ΔEx5’/ΔEx5’) (Figure S8B). Levels of flTCF7L2 had not changed relative to +/+ mice, however dnTCF7L2 could not be detected (Figure 6G and 6H), suggesting that the brain-specific dnTCF7L2 is indeed translated from these exon 5’-containing transcripts. Overall, dnTCF7L2 level in the midbrain of ΔEx5’/+ mice was comparable to that of +/+ mice, indicative of a compensatory mechanism. Importantly, we found severe USV abnormalities in Δ 5’/ Ex5’ but not ΔEx5’/+ and +/+ pups (Figure 6I), indicating that brain-specific dnTCF7L2 is required for mouse USV production.

### Decrease of synaptic transmission in lateral PAG with loss of TCF7L2

To understand the molecular basis underlying TCF7L2 loss-associated mouse USV impairment, we performed RNA profiling of the PAG region in P7 *exon11 fl*/*fl*;*Vglut2*-*Cre*/+ mice, in which both flTCF7L2 and dnTCF7L2 are depleted in *Vglut2*-positive neurons (Figure S7A). Gene Set Enrichment Analysis (GSEA) revealed that several biological pathways were altered in *exon11 fl*/*fl*;*Vglut2*-*Cre*/+ pups relative to *exon11 fl*/*fl* pups (Figure 6J). Specifically, gene sets involved in neuron projection membrane (normalized enriched score, NES = -2.11, *p* < 0.001), potassium channel activity (NES = -1.92, *p* < 0.001), and synaptic vesicle priming (NES = -1.97, *p* < 0.01) were significantly downregulated in the cKO PAG region, suggesting impaired neuronal function upon depletion of TCF7L2 in the *Vglut2*-positive neurons (Figure 6K). Amongst upregulated gene programs we found genes involved in SMAD signaling (NES = 1.92, *p* < 0.001) and sugar membrane transporter activity (NES = 1.91, *p* < 0.001), suggesting that both BMP/SMAD signaling and glucose metabolism is altered in the cKO PAG (Figure 6K).

To investigate whether loss of TCF7L2 causes PAG neuronal dysfunction, we performed whole cell current clamp of the lateral PAG (LPAG) region in acute slices from one-month old Δ2/+ and +/+ mice (Tschida et al., 2019). Overall, haploinsufficiency of *Tcf7l2* did not significantly change neuron excitability nor spike features in LPAG neurons (Figure S9). However, the miniature excitatory postsynaptic current (mEPSC) frequency and miniature inhibitory postsynaptic current (mIPSC) amplitude were significantly Δ2/+ LPAG neurons relative to +/+ animals (Figure 6L and 6M). Taken together, our data suggests that haploinsufficiency of *Tcf7l2* alters neuronal gene expression profiling in a way that impairs LPAG neuron synaptic transmission.

### Mice carrying disease-associated mutations produce less USVs and haploinsufficiency of *Tcf7l2* does not lead to ASD-like behaviors in mouse

Specific *TCF7L2* mutations have been associated with development disorders (DD), autism spectrum disorder (ASD), and language delay (Deciphering Developmental Disorders, 2015; Dias et al., 2021; Iossifov et al., 2014; Stessman et al., 2017) (Figure 7A, S10, and Table S1). To investigate whether disease-associated *TCF7L2* mutations impair mouse USV production, we selected two, c.932+1G>A and c.1150C>T (R384X), to generate knock-in mouse lines (Figure 7B and 7C). The c.932+1G>A mutation alters the 5’ splicing site (GT) of *TCF7L2* exon 9 to AT, potentially preventing correct splicing from exon 9 to exon 10, the latter of which encodes part of the HMG domain. The c.1150C>T (R384X) mutation introduces a premature termination codon in *TCF7L2* exon 11 (which also encodes the HMG domain) (Figure 7A), and we therefore speculated that both mutant transcripts would be targeted by nonsense mediated decay for degradation. Indeed, we found that midbrain expression of both flTCF7L2 and dnTCF7L2 in mice heterozygous for the two disease-associated mutations (A/+ and T/+) was reduced to half of that of wildtype littermates (Figure 7D and 7E), suggesting that the two disease-associated mutations are loss-of-function alleles.

**Figure 7.**
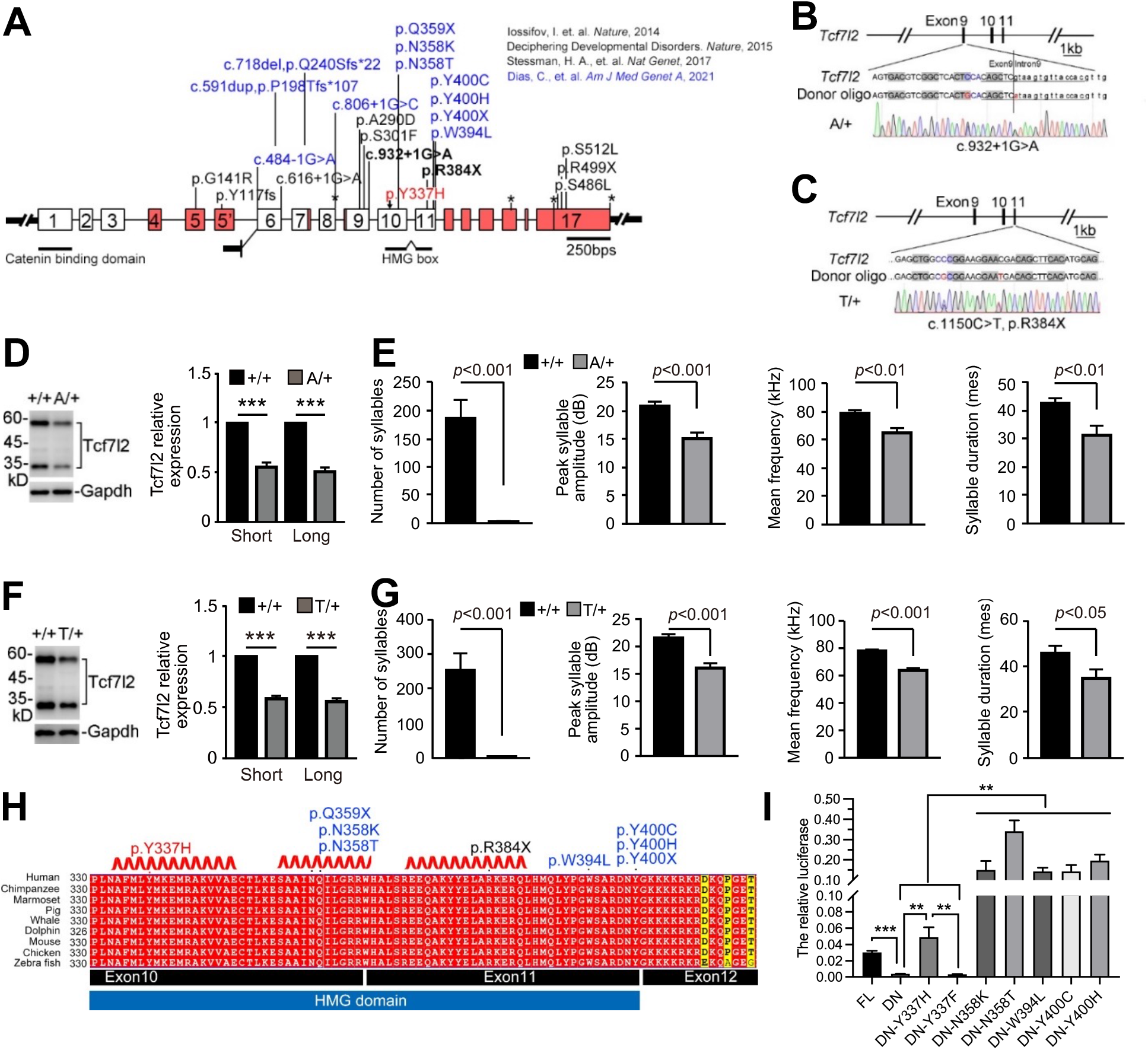
Disease-associated mutations impairs mouse USVs by relieving the transcriptional repression of TCF7L2. (A) Gene structure of *TCF7L2* and human mutations found in patients (PMID: 25363768, 25533962, 28191889, and 34003604). (B-E) Generation of *Tcf7l2* knockin mouse lines carrying disease-associated mutations. TCF7L2 expression in P7 midbrains with indicated genotypes (D and E). (F and G) The syllable numbers, mean frequency, syllable duration, and peak syllable amplitude measured at P7 with indicated genotypes. (H) Protein sequence alignment of TCF7L2 HMG domain. Human TCF7L2 (NM_001198528) were employed for positioning. Disease-associated mutations include: 1) our ENU-induced Y337H mutation colored in red; 2) mutations described in previous studies are colored in black (PMID: 25363768) and blue (PMID:34003604), respectively. (I) Transcriptional repression of dnTCF7L2 (DN), which lacks of CBD (beta-catenin binding domain), was relieved by the disease-associated mutations shown in H. Previously reported TOPFlash luciferase reporter (PMID: 9065401) was employed for the measurement. FL, full-length TCF7L2. Renilla luciferase activity was used for normalization. The value are presented as mean ± SEM. N.S., no significant difference, ** *p*<0.01, *** *p*<0.001, t-test or ANOVA, SPSS. In D, F, and I, n=3; in E and G, +/+ (n=10), A/+ (n=8), and T/+ (n=10). See also Table S1, Figure S10-S12.

Similar to the C/+ and Δ2/+ mice, the A/+ and T/+ pups generated fewer syllable numbers, lower peak amplitude, reduced mean frequency, and shorter duration compared to control mice (Figure 7F and 7G). Therefore, we conclude that the two *TCF7L2* disease-associated mutations are likely loss-of-function mutations and impair mouse USV production in a haploinsufficient manner, like the ENU-induced Y337H and Δ2 KO alleles.

To examine whether haploinsufficiency of *Tcf7l2* leads to ASD-like behaviors, we performed three-chamber test in C/+ and Δ 2/+ spent significantly more sniffing time towards their conspecific than object (Figure S11A-S11D) and social preference indexes of C/+ and Δ2/+ relative to control mice were comparable, suggesting that *Tcf7l2* haploinsufficiency in and of itself does not impair social ability. When we next assessed repetitive behaviors through a marble burying test and estimated self-grooming, we found that C/+ and Δ + mice buried similar percentage of marbles and spent similar self-grooming time relative to +/+ mice (Figure S11E and S11F), hence *Tcf7l2* haploinsufficiency is not associated with ASD-like behaviors in mouse.

### The disease-associated non-synonymous mutations in HMG domain relieve transcriptional repression of dnTCF7L2

Besides our ENU-induced Y337H mutation, many disease-associated mutations are found in the HMG domain, which is highly conserved from fish to human (Figure 7H and Figure S10). These mutations include three stop-gain mutations (Q359X, R384X, and Y400X) and five non-synonymous mutations (N358K, N358T, W394L, Y400H, and Y400C). Here, we found that knock-in of the R384X mutation lead to severe USV abnormalities in mouse (Figure 7F and 7G), suggesting that the other two stop-gain mutations are likely dysfunctional mutations. Indeed, patients with the Q359X and Y400X mutations displayed severe speech delay, as well as that with the 5 non-synonymous mutations (Table S1).

Because both flTCF7L2 and dnTCF7L2, the latter of which lacks the transcriptionally active domain, are required for vocal production (Figure 6), we hypothesized that a transcriptional repression mechanism of TCF7L2 is crucial for the complex trait. To this end, we expressed flTCF7L2 and dnTCF7L2 in HEK293 cells and measured their transcriptional activities with an established TCF reporter system (TOPFlash) (Korinek et al., 1997). Consistent with previous studies (Korinek et al., 1997; Vacik et al., 2011), the transcriptional activation and repression was achieved by expression of flTCF7L2 (FL) and dnTCF7L2 (DN), respectively (Figure 7I). However, we found that the repression effect was abolished by Y337H (DN-Y337H) but restored by Y337F (DN-Y337F), indicating that the DNA-binding ability of TCF7L2 is required for the effect. Like Y337H, five disease-associated non-synonymous mutations (DN-N358K, DN-N358T, DN-W394L, DN-Y400H, and DN-Y400C) relieved the transcriptional repression of dnTCF7L2. Therefore, our data is most consistent with a model in which the disease-associated mutations in HMG domain, like Y337H, relieve the transcriptional repression of TCF7L2, which in turn leads to vocal abnormalities in patients.

## Discussion

Here, through the use of a battery of genetic mouse models, we show that TCF7L2 acts as a molecular switch in midbrain PAG *Vglut2*-positive neurons to control mouse USV production and syllable complexity, probably through a transcriptional repression mechanism. Previously, transcriptional repression mechanisms and Wnt/β signaling activation independent roles of TCF7L2 have been documented during oligodendrocyte differentiation and brain development (Hammond et al., 2015; Vacik and Lemke, 2011; Vacik et al., 2011; Ye et al., 2009; Zhang et al., 2021; Zhao et al., 2016). In addition, transcriptional repression of BMP4/SMAD signaling by TCF7L2 has been linked to oligodendrocyte differentiation (Zhang et al., 2021). In agreement with these findings, we observed that expressions of genes involved in BMP/SMAD but not Wnt/ β-catenin signaling are upregulated in *Tcf7l2* cKO PAG (Figure 6K). In addition, TCF7L2-mediated vocal control is independent of its β-catenin-binding domain (Figure 6) but dependent of its DNA binding ability (Figure 2). Given that dnTCF7L2 inhibits the Wnt/β-catenin signaling cascade (Cadigan and Waterman, 2012; Hoppler and Kavanagh, 2007; Vacik and Lemke, 2011; Vacik et al., 2011), and is required for the vocal control, we argue that a transcriptional repression mechanism mediated by TCF7L2 is crucial for control of the complex vocalization trait. Indeed, our data how that like Y337H, patient mutations associated with speech delay relieve the transcriptional repression of TCF7L2 (Figure 7I).

As we demonstrate that loss of TCF7L2 and the Y337H mutation in the HMG domain result in vocalization abnormalities in mice, it is important to note that patients carrying *de novo* genetic variants, including loss-of-function mutations and non-synonymous mutations in HMG domain similar to what we reported here, display severe speech delay, developmental disorder, and ASD (Deciphering Developmental Disorders, 2015; Dias et al., 2021; Iossifov et al., 2014; Stessman et al., 2017). Knock-in mice with such disease-associated mutations displays severe USV abnormalities, indicating that the cellular and molecular mechanisms we identify here are most likely shared by mouse and human. Consistent with less ASD penetration in the patients (Dias et al., 2021), we did not detect ASD-like phenotypes, including social disability and repetitive behaviors in the *Tcf7l2* mutant mice (Figure S11).

Given that PAG functions as an organizing node for mammal vocalization in the brain (Jurgens and Hage, 2007; Nieder and Mooney, 2020; Tschida et al., 2019), the impaired synaptic transmissions in LPAG we detected here may be the consequence of TCF7L2 loss of function, thereby explaining the mechanisms of vocalization deficits (Figure 6L and 6M). In addition to PAG’s role as a central vocalization node, *Esr1*-positive neurons in the ventromedial hypothalamus and preoptic area (POA) and *Vgat*-positive inhibitory neurons in amygdala (Amg) act upstream of PAG-USV neurons to contribute to mouse USV production and persistence (Chen et al., 2021; Gao et al., 2019; Karigo et al., 2021; Michael et al., 2020). We demonstrate that expression of TCF7L2 in *Vglut2*-positive neurons is essential for mouse USV production, excluding the possibility that expression of TCF7L2 in *Vgat*-positive neurons in POA, Amg, and PAG contribute to the processes. By removing *Tcf7l2* in *Esr1*-positive and *ChAT*-positive neurons, we further exclude the possibilities that *Esr-1*-positive neurons in VMH and POA and *ChAT*-positive neurons in medial habenula and spinal cord motor neurons participate in the processes (Figure S12). Given that *TCF7L2* has been associated with metabolic disorders (Grant et al., 2006) and glucose metabolism is affected in PAG by loss of TCF7L2 in *Vglut2*-positive neurons (Figure 6J and 6K), metabolic status may influence PAG-USV neuron activities and contribute to mammal vocal production and performance. By nanopore sequencing, we demonstrate that *Tcf7l2* undergoes multiple alternative splicing, including at least 10 different exon5’ usages for transcripts encoding dnTCF7L2, in midbrain (Figure 6C). In future work, it will be intriguing to investigate *TCF7L2* alternative splicing patterns across different species to syllable complexity and human language evolution.

## Material and methods

### Animals

Experimental procedures were approved by AAALAC (Association for Assessment and Accreditation of Laboratory Animal Care International) and the IACUC (Institutional Animal Care and Use Committee) of Tsinghua University. All the mice described in this research were C57BL/6J background and housed in room at 22-26°C in 12/12-hour light/dark cycle with sterile pellet food and water ad libitum. The *Vglut2*-cre mouse (# 016963, Jackson Laboratory) was gifted from Dr. Minmin Luo at NIBS. *Olig3*-cre (Vue et al., 2009) was kindly provided by Dr. Songhai Shi; *Esr1*-cre (#017911, Jax) was gifted from Dr. Xiaohong Xu; *Tcf7l2* exon11 floxed mouse was gifted from Dr. Richard Lu; *Nestin*-cre (#003771, Jax), *ChAT*-cre (#006410, Jax), and *Tcf7l2* exon1 floxed mouse (#031436, Jax) were imported from The Jackson Laboratory.

### Forward genetics screening and mutant candidate identification

Male C57BL/6J mice were treated with ENU (80-110 mg/kg B. W.) for three consecutive weeks by intraperitoneal injection at the age of 12 weeks according previously reported (Chen et al., 2020). The ENU-treated males were crossed with C57BL/6J females and the resulting G1 pups were subjected to USV measurement at P5 with maternal deprivation in a 5-minute interval by a commercial USV detector (Med Associates Inc).

The USV impaired G1 mice (≤5 in the 5-minute interval we measured) were crossed to C57BL/6J and the recurrence of USV impairment in G2 and G3 offsprings was used to establish the family pedigree and the genomic DNA was applied for whole exome sequencing. Mouse genomic DNA from USV affected and unaffected mice in the family bp fragments and captured by SeqCap EZ probe pool (Roche). The libraries were sequenced through HiSeq and the reads were aligned to GRCm38 mouse reference genome. The mutant candidates were selected by previously described criteria (Chen et al., 2020).

### Knockout and FLEx (fx) conditional knockin mouse generation by CRISPR/Cas9 technology

All the sgRNAs were designed by Cas-Designer (http://www.rgenome.net/cas-designer/). Cas9 mRNA and sgRNAs were synthetized and purified by HiScribe™ T7 High Yield RNA Synthesis Kit (E2040S, NEB) and Purification of Synthesized RNA (E2040, NEB), respectively. Mixed Cas9 mRNA, sgRNAs, and/or donor DNA were injected into C57BL/6J mouse zygotes at the pronuclei stage. The injected eggs were transferred to into uterus of pseudopregnant ICR females. For generation of *Foxp2* Δ allele, the sgRNA: CCATACGAAGGCGACATTCAGAC; for *Tcf7l2* Δ2 allele, the sgRNA: TATGAAAGAGATGAGAGCGA; for *Tcf7l2 fx*, the sgRNAs: CCAGTGATTGAAACTGACAC, CCAGTGTCAGTTTCAATCAC, GATTCCACCCAGGCAGAAAA, TAACGGGTTTGATTCCACCC; for *Tcf7l2* ΔEx5’, the sgRNAs: ATGAATGATCGGGTTCTTTG, GAAGAACTGAGACAAAATGT, CCACAGAGGACTGCTACCAT, CCTATGGTAGCAGTCCTCTG. The donor DNA for *Tcf7l2 fx* mouse include the inverted exon9 and exon10 and their surrounding sequences flanked by two pairs of incompatible lox sites (loxP and lox2722) and homology arms (∼3kb).

### Immunofluorescence, RNA in situ hybridization, and western blot

Mice was anesthetized and transcardially perfused with cold PBS and 4% PFA. Brains were post fixed and dehydrated with 30% sucrose and sectioned to 20 μm. The sections were blocked and permeabilized with 3% BSA and 0.3% Triton X-100 in PBS, and then incubated with the primary antibody overnight at 4°C. Sections were washed with PBST three times and incubated with the secondary antibody. The antibodies: rabbit anti-TCF7L2 (Cell Signaling Technology, #2569, 1:200), mouse anti-HA (Abcam, ab18181, 1:500), donkey anti-mouse IgG secondary antibody (Alexa Fluor 555, Thermo Fisher Scientific, A31570, 1:1000), donkey anti-mouse IgG secondary antibody (Alexa Fluor 488, Thermo Fisher Scientific, A32766, 1:1000). RNA in situ hybridization was performed according to RNAscope manual. For western blot, tissues were homogenized in RIPA buffer (50 mM Tris-HCl, pH 8.0, 150 mM NaCl, 0.25% sodium deoxycholate, 0.1% SDS, and 1% NP-40) supplemented with complete protease inhibitor mixture (Roche). Lysates were centrifuged at 12000 x g for 10 minutes at 4°C. The membranes were incubated with the primary antibodies overnight at 4°C, then washed with PBST and incubated with HRP-conjugated secondary antibodies. Intensities of blot bands were calculated by Image J.

### Protein expression and purification and electrophoresis mobility shift assay (EMSA)

The coding sequences of wild type and mutant HMG domain were cloned into pMAL-cRI plasmid with MBP (Maltose Binding Protein) tag. The expression of recombinant HMG proteins was induced by 0.3 mM IPTG in BL21 at 30°C. Cells were lysed by ultrasonic homogenizer. After centrifugation, the recombinant proteins were purified by amylose resin (#E8022S, New England Biolabs) according the manual instructions. EMSA was performed and HMG-binding sequences (5’-tagacgtagggcaccctttgaagctctccctcga) were synthesized as previously described (Giese et al., 1991). Briefly, biotin-labeled probe was incubated with different HMG proteins in EMSA buffer (50 mM NaCl, 20 mM Tris-HCl, 1 mM DTT, 1 mM MgCl_2_, 5% Glycerol) for 1 hour at 25°C. Then, the samples were separated with 6% native PAGE, and transferred to nylon membranes. After UV crosslinking, the membrane was incubated with HRP-conjugated Streptavidin. Finally, the signal was visualized by GE imaging system.

### Luciferase assay

The plasmid of TOPflash, pCMV-Flag-β-catenin, and pCMV-Renilla together with each dnTCF7L2 point mutation (DN-Y337H, DN-Y337F, DN-N358K, DN-N358T, DN-W394L, DN-Y400H, and DN-Y400C) were transfected into HEK293T cells in six-well plates. Cell lysis were harvested 12 hours after transfection for luciferase analysis according to the manual instruction (#RG027, Beyotime). The value of luciferase was measured by VARIOSKAN FLASH (Thermo).

### Reverse Transcription PCR (RT-PCR), real-time PCR, RNA-Seq, and nanopore sequencing

For RT-PCR, total RNA was isolated by TRIzol reagent (Thermo Fisher Scientific). RNA was treated with DNase and cDNA was synthesized by M-MLV Reverse Transcriptase (Promega). For real-time PCR, the reaction was performed with 2 x Hieff qPCR SYBR Green Master Mix (Yeasen) and detected by LightCycle 480 Real-Time PCR machine (Roche). For RNA-seq, RNA sample library preparation was performed according to TruSeq mRNA-seq Stranded v2 Kit (Illumina) manual instruction. The library was sequenced using Illumina Hiseq platform. Reads were aligned to GRCm38 mouse reference. HTSeq was used to quantify the aligned reads (Anders et al., 2015). Reads normalization and differential gene expression analysis were performed using DESeq2 (Love et al., 2014). Normalized reads were subjected to Gene Set Enrichment Analysis (GSEA) (Subramanian et al., 2005). For nanopore sequencing, RNA library was constructed following manual instruction (Oxford Nanopore Technologies) and the sequencing was conducted using Oxford Nanopore Technology’s MinION. Long reads were aligned to GRCm38 mouse reference using minimap2 (Li, 2018). Isoform definition and quantification were achieved by FLAIR (Tang et al., 2020). Reads were visualized by Integrative Genomics Viewer (IGV) (Robinson et al., 2011).

### AAV preparation and injection

AAV2/9 Syn-Cre-mCherry and Syn-mCherry were purchased from OBiO (Shanghai). After born within 6 h, the mice were anesthetized with ice and the head was fixed by plasticine in stereotaxic apparatus. AAV virus (60 nl) were infused by glass micropipettes at a rate of 50 nl/min bilaterally in PAG region (-0.8 mm AP from Lambda; ± 0.35 mm ML; -1.85 mm DV). The injected mice were put on warm blanket to recovery 1 hour at 37°C.

### Electrophysiology

Electrophysiological recording was performed with MultiClamp 700B amplifier (Molecular Devices, USA) as previously described (Li et al., 2017). Mice (P25-P30) were anesthetized and transcardially perfused with ice-cold oxygenated cutting aCSF, containing (in mM) 125 sucrose, 1.25 NaH_2_PO_4_, 2 CaCl_2_, 3 KCl, 2 MgSO_4_, 26 NaHCO_3_, 1.3 sodium ascorbate, and 0.6 sodium pyruvate. PAG slices (300 μm) were prepared and recovered in normal aCSF, containing (in mM): 125 NaCl, 1.25 NaH_2_PO_4_, 2 CaC_l2_, 3 KCl, 2 MgSO_4_, 26 NaHCO_3_, 1.3 sodium ascorbate, and 0.6 sodium pyruvate, for 25 minutes at 34°C, and then maintained at room temperature for 1 hour before recording. For mEPSC, slices were transferred into recording chamber perfused with aCSF in the presence of 1 μM tetrodotoxin and 50 μM picrotoxin. Recording micropipettes were prepared by Sutter P97 puller with a resistance of 3-5 MΩ and filled with internal solution, containing (in mM): 130 potassium-gluconate, 20 KCl, 10 HEPES, 10 Na_2_HPO_4_, 4 Mg_2_ATP, 0.3 Na_2_GTP, and 0.2 mM EGTA (pH 7.35, 299 mOsm). For mIPSC, the slices were bathed in aCSF with 10 μM CNQX and 50 μM AP5 and recording internal solution containing (in mM): 60 CsCl, 2 MgCl_2_, 10 HEPES, 10 Na_2_HPO_4_, 4 Mg_2_ATP, 0.3 Na_2_GTP, and 0.2 mM EGTA (pH 7.35, 299 mOsm). For neuron excitability measurement, action potential firing was recorded under current clamp mode with pipette filled with the internal solution used for mEPSC. Neuron was injected with currents from 0 to 300 pA with an increment of 50 pA.

### Mouse behavior analysis

#### Open field

Mice were placed in the box (50 × 50 × 50 cm) and freely explored for 30 minutes. Total travel distance and time spent in the center area were calculated by TopScan (CleverSys Inc.) behavior analysis software.

#### Rotarod performance test

Mice were put on the accelerating rotarod with speed from 4 to 40 rpm in 5-minute and three consecutive trials per day for 3 days. A 2-minute break was set in between each trial. The fall time from the spindle was auto-calculated by the system (Med Associates Inc.).

#### Ultrasonic vocalization (USV) measurement

For pup vocalization, pup was isolated from their littermates and mom and placed into a glass cup in a soundproof box. USVs were recorded by ultrasound microphone (Avisoft CM16/CMPA) for 5 minutes. For adult mouse USV measurement, 3-month-old males were single caged for 5 days before the test. After the tested male was habituated in the soundproof box for 5 minutes, live or anesthetized virgin female was placed into the box and USVs were recorded. For analysis of USVs, syllable numbers, peak syllable amplitude, mean frequency, and syllable duration were measured by MUPET (Van Segbroeck et al., 2017). Syllable repertoire composition was classified into 4 categories (simple syllables “s”, two notes syllables downward “d” and upward “u”, multiple syllables “m”) by Mouse Song Analyzed v1.3 (Chabout et al., 2017). Syllable transitions of conditional probability for each context were analyzed by a customized script generated in Microsoft Excel (Chabout et al., 2017).

#### Three-chamber social ability test

A rectangular box (60 length × 40 width × 25 height, cm) consisted of three chambers (20 m) side by side. The tested mouse was allowed to habituate the chambers for 5 minutes and then placed in the center chamber. A stranger mouse (S) was placed in a container in one side of the chamber and an object (O, another container) was placed in the other side. Then the tested mouse was allowed to explore S and O for 10 minutes and the sniffing time toward S and O was calculated manually. Social preference index was calculated as previously described (Dong et al., 2020).

#### Female preference test

Female was placed in the three chamber as described above and allowed to explore the chamber for 5 minutes, and then placed in the center chamber. Wildtype (+/+) mouse was placed in a container in one side chamber and C/+ or Δ mutant mouse was placed in the other side. Then the tested female was allowed to explore the chamber freely for 20 minutes and the sniffing time toward each mouse was calculated manually.

#### Marbles buried test

Fresh bedding (5 cm) was filled in a standard mouse cage with 20 dark glass marbles (1.5 cm diameter), which were placed on the bedding in a symmetrical 4 × 5 pattern. Each tested mouse was allowed to explore freely for 30 minutes. The number of buried marbles (> 2/3 height of marble covered with bedding) was counted.

#### Self-grooming test

The tested mouse was placed in a standard mouse cage with fresh bedding and videotaped for 20 minutes. The time of self-grooming behaviors (including rubbing/scratching head, face, and other body parts) were manually accounted.

---

**Figure S1.**
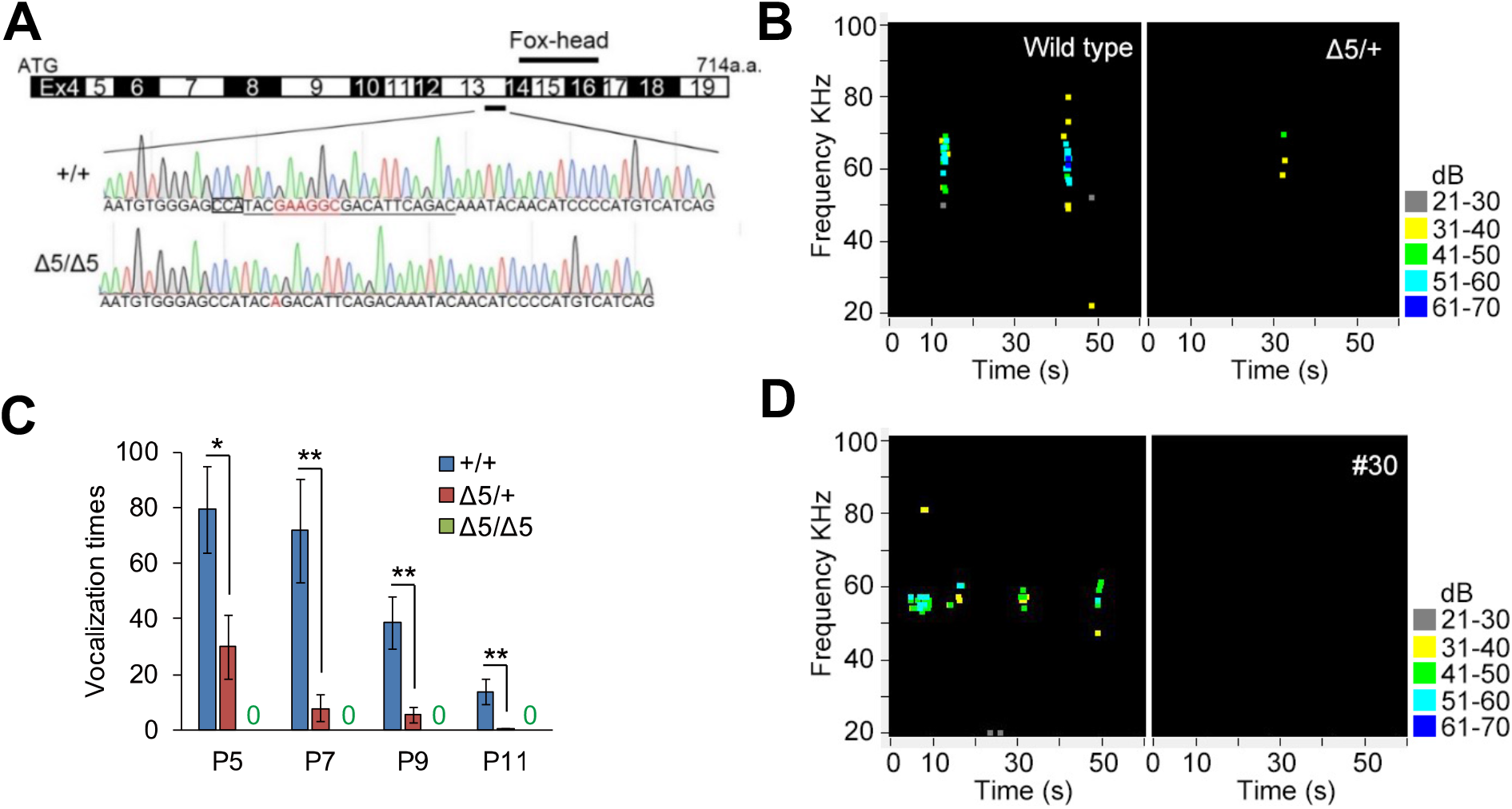
*Foxp2* KO impairs mouse USV production. (A) A CRISPR/Cas9-based *Foxp2* KO mouse. The gRNA designed for the KO is located in *Foxp2* exon13 upstream of the exons coding Fox-head domain. A 5- nucleotide deletion (Δ5) generated by CRISPR/Cas9-mediated editing and the mouse homozygous for the deletion (Δ5/Δ5) was applied for the Sanger sequencing. The gRNA sequences were underlined and PAM (NGG) site was boxed. (B) Pup USVs of the wildtype and heterozygous KO (Δ5/+) pups at P5 were measured by a USV detector (Med Associates Inc.). A representative USV impairment of Δ5/+ mouse P5 was shown. (C) Both Δ5/+ and Δ5/Δ5 mice displayed the USV impairment at P5, 7, 9, and 11. (D) Family members from #30 displayed USV impairment at P5. In C, data are presented as mean ± SEM, +/+, n=15; Δ5/+, n=14; Δ5/Δ5, n=3, * *p*<0.05, ** *p*<0.01, t-test, SPSS.

**Figure S2.**
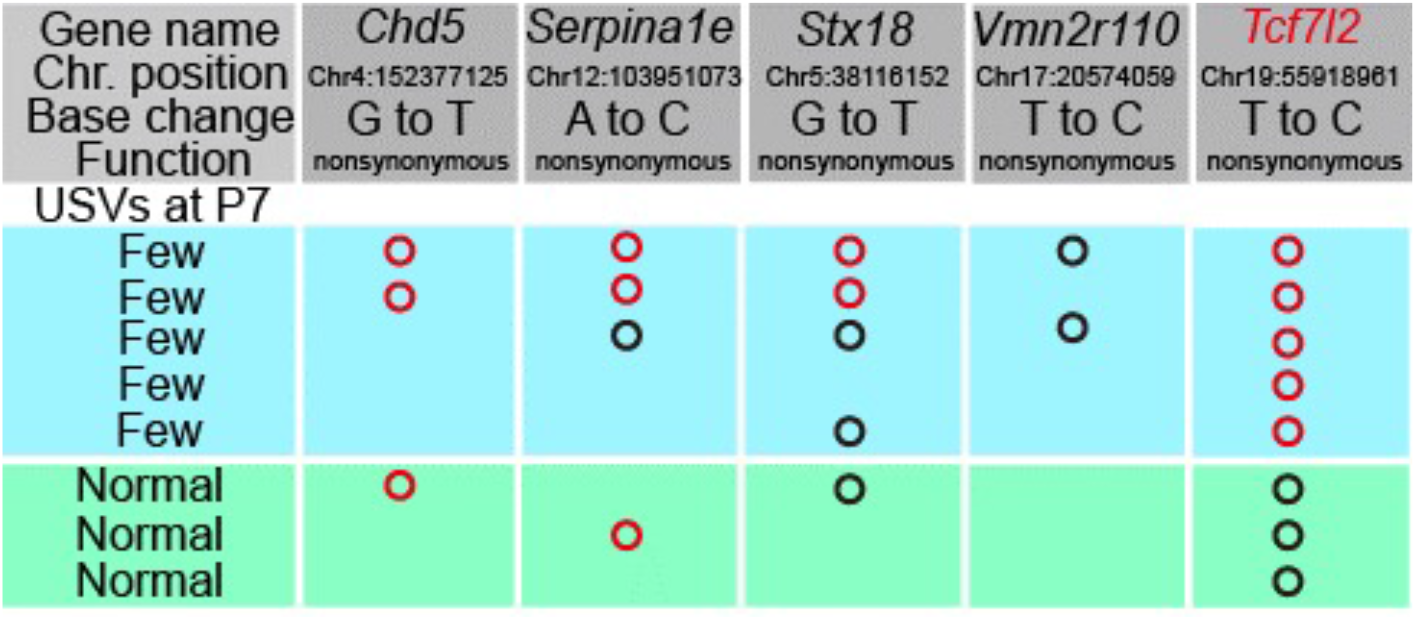
Identification of an ENU-induced *Tcf7l2* mutation that impairs pup USVs. Some of nonsynonymous mutations identified by whole-exome sequencing are shown, which were tested in 5 affected (Few USVs) and 3 unaffected (Normal USVs) mice in the #30 family. The *Tcf7l2* mutation was co-segregated with the USV impairment at P7.

**Figure S3.**
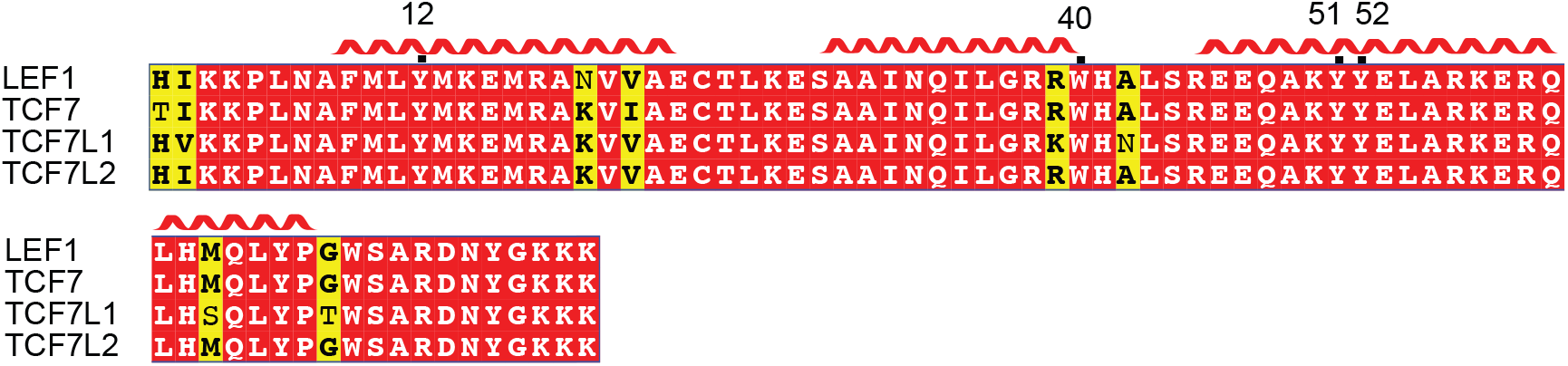
Aromatic residues Y12, W40, Y51, and Y52 in LEF1 HMG box are shared by the TCF/LEF family members. An alignment of HMG boxes of 4 human TCF/LEF family members and Y12, W40, Y51, and Y52 are highlighted.

**Figure S4.**
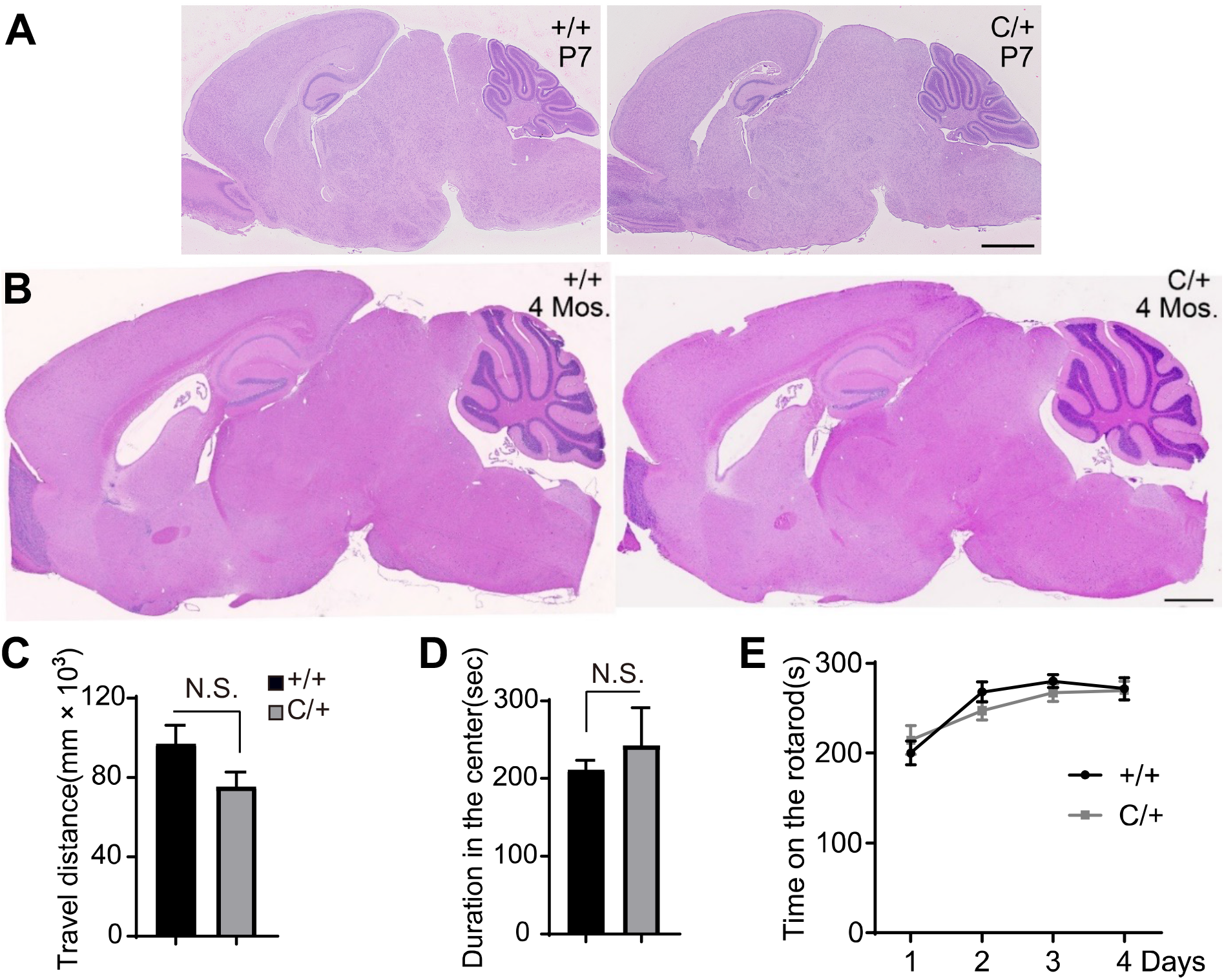
Normal brain morphologies and motor abilities shown in the ENU-induced mutant animals. (A and B) Hematoxylin-eosin staining of wildtype (+/+) and the ENU-induced mutant (C/+) brains at P7 (A) and 4 months of age (B). Scale bar, 1mm. (C-E) The mice with indicated genotypes were applied for Open field (C and D) and Rotarod tests (E). The value are presented as mean ± SEM. In C and D, +/+, n=10; C/+, n=9; in E, +/+, n=12; C/+, n=12. Mice, male, 3 months of age. N.S., no significant difference, t-test, SPSS.

**Figure S5.**
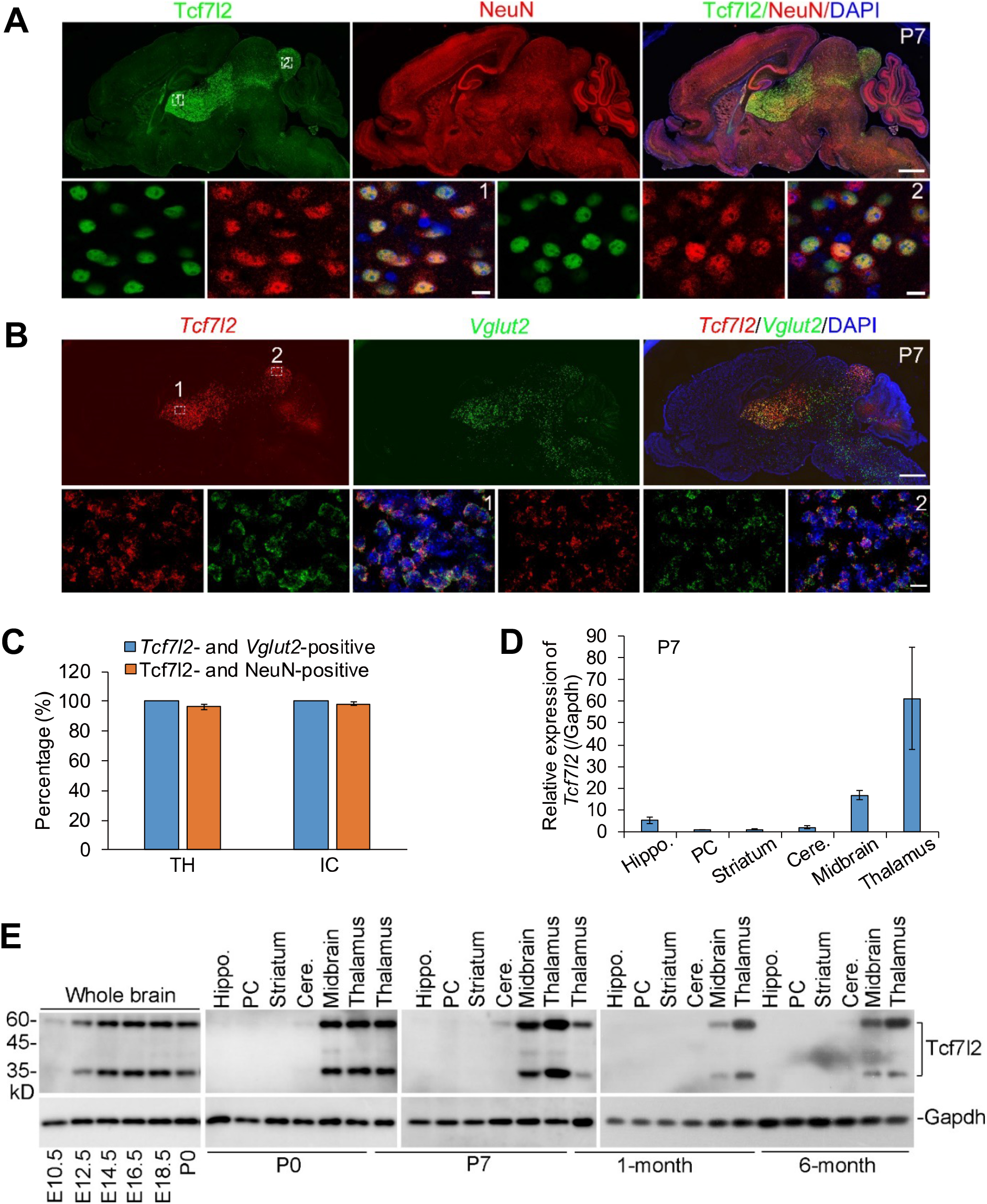
Expression pattern of TCF7L2 protein and*Tcf7l2* mRNA in mouse brain. (A and B) Immunostaining of TCF7L2 and NeuN (A) and *in situ* hybridization of *Tcf7l2* and *Vglut2* (B) in a mouse sagittal section at P7. Enlarged region 1 (Thalamus, TH) and 2 (Inferior colliculus, IC) were shown (Lower). (C) Statistical analysis of double positive neurons in region 1 (TH) and 2 (IC) at P7. (D) The expression of *Tcf7l2* in various brain tissues detected by the real-time PCR at P7. (E) Expression pattern of TCF7L2 was examined by western blot. GAPDH, loading control. In C and D, the value are presented as mean ± SEM (n=3). In A and B, scale bar, 1mm (low magnification) and 20 μm (high magnification).

**Figure S6.**
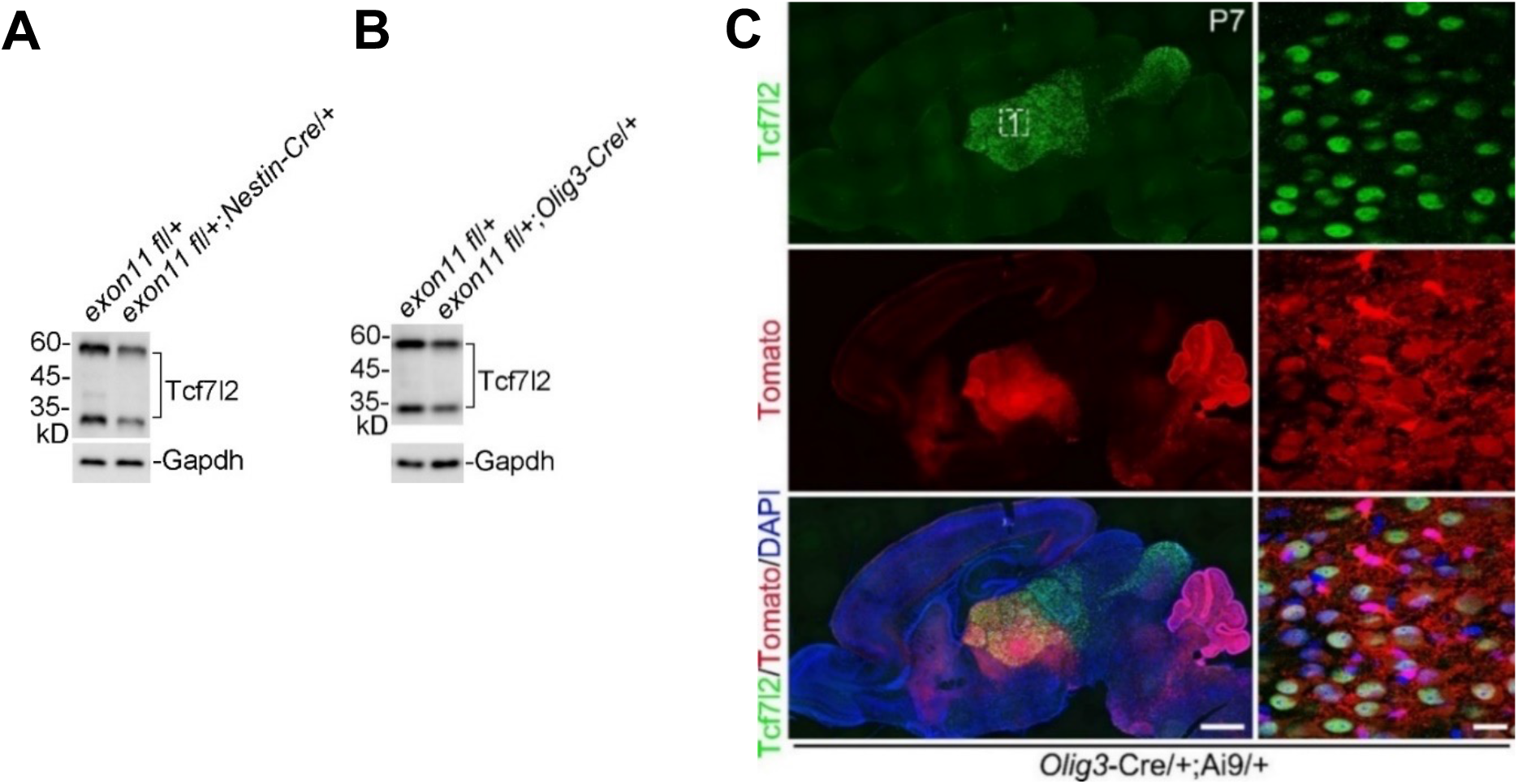
Expression of TCF7L2 in midbrains in the cKO mice and *Oligo3*-*Cre* and TCF7L2 expression pattern at P7. (A and B) The expression of TCF7L2 in the indicated cKO mice by western blot. GAPDH, loading control. (C) *Oligo3* expression was reported by Ai9 reporter mouse (PMID: 20023653). TCF7L2 expression was measured by immunostaining. Enlarged region 1 (Thalamus) was shown in the right. Scale bar, 1mm (left, low magnification) and 20 μm (right, high magnification).

**Figure S7.**
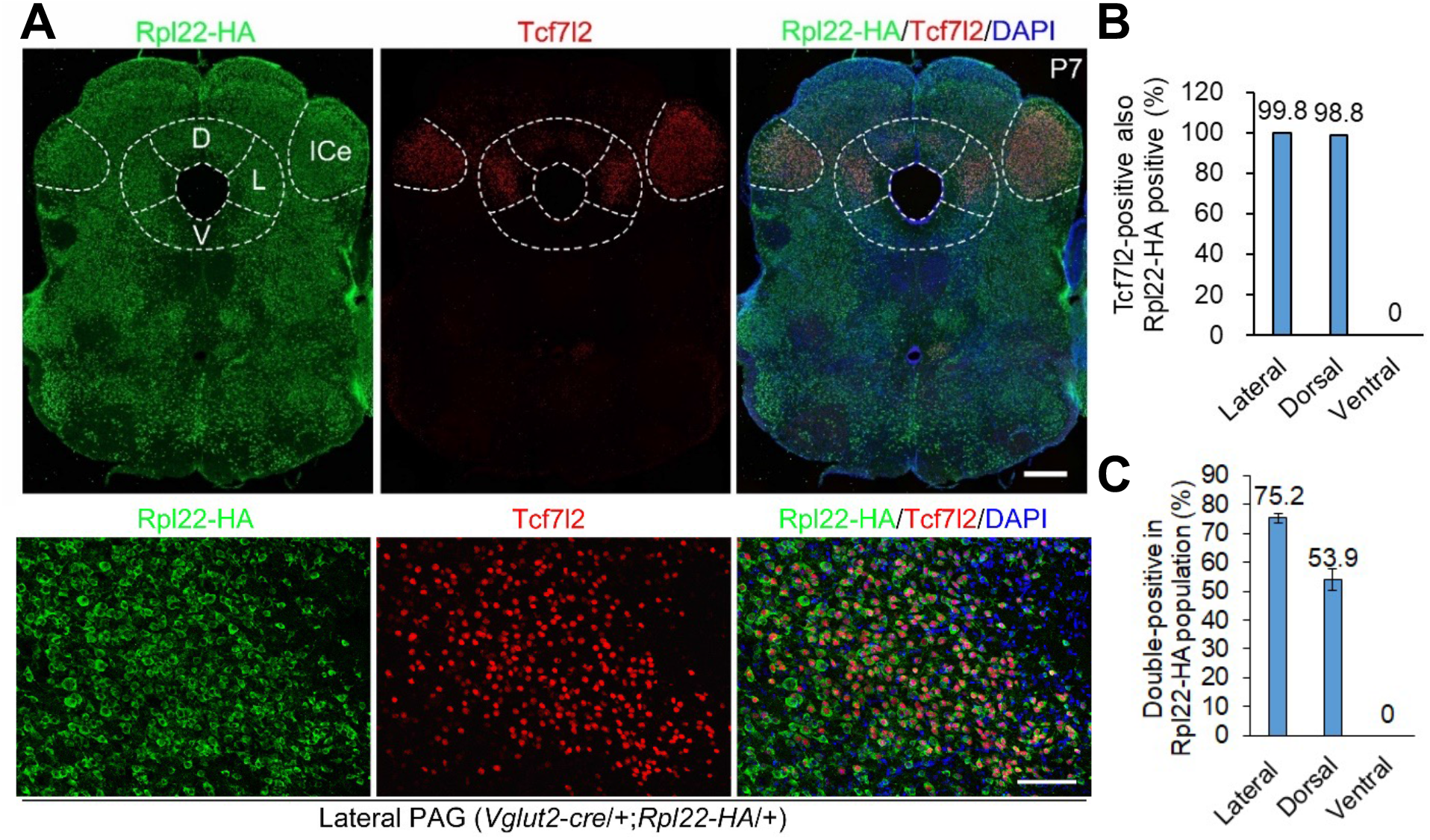
Expression of TCF7L2 in PAG at P7. (A) The RiboTag mice (PMID: 19666516) were crossed to *Vglut2-Cre*/+ mice to label *Vglut2-*positive neurons in midbrain PAG. Neurons with TCF7L2 immunoreactive nuclear signals were positive for RPL22-HA cytosolic signals in PAG at P7. Scale bar, 500 μm (top) and 50 μm (bottom). L, lateral; D, dorsal; V, ventral; ICe, external cortex of the inferior colliculus. PAG and ICe was shaped by dashed lines. In B and C, the value are presented as mean ± SEM.

**Figure S8.**
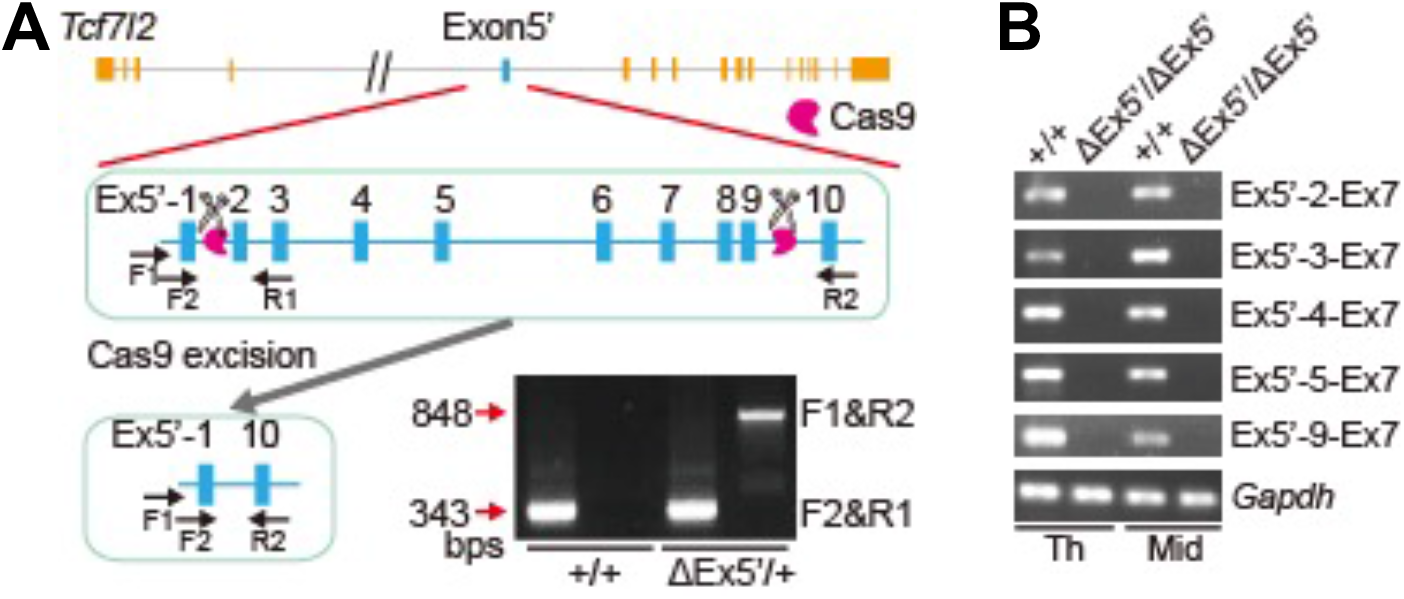
Generation of *Tcf7l2* exon5’ KO mouse. (A) A ∼6kb DNA fragment containing exon5’-2 to -9 was removed by CRISPR/Cas9- based KO. We employed two sgRNAs (labeled as scissors). Ten alternative exon5’ of *Tcf7l2* are represented as blue rectangles. Primers for genotyping are labeled (F1, R1, F2, and R2). An additional 848bp band was only amplified by genomic DNA PCR in mouse heterozygous for exon5’ KO (ΔEx5’/+) but not +/+. (B) Expression of exon5’-containing transcripts detected in +/+ but not ΔEx5’/ΔEx5’ mouse.

**Figure S9.**
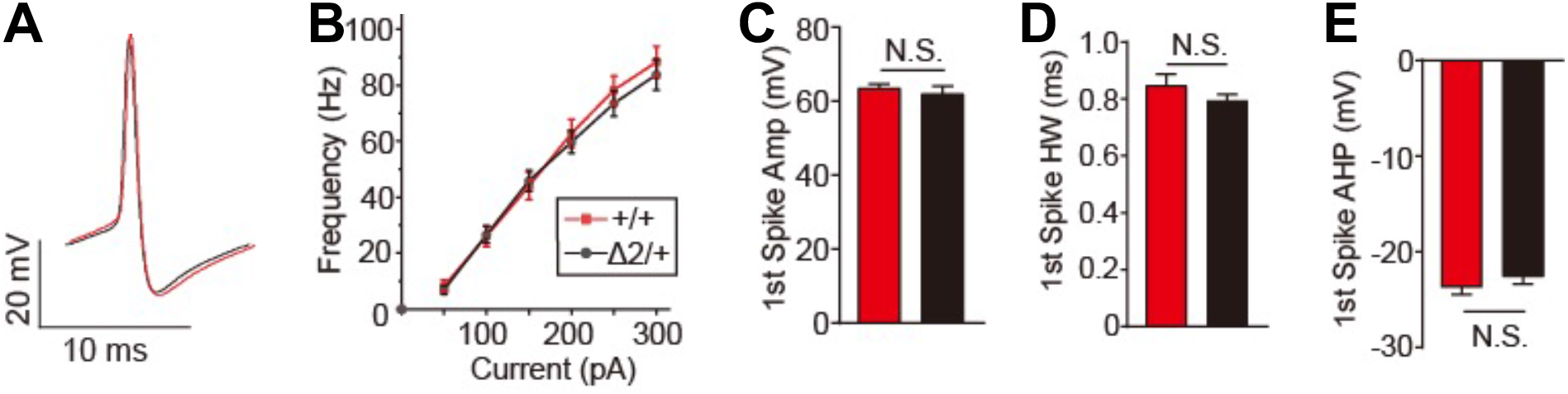
Haploinsufficiency of *Tcf7l2* does not affect neuron intrinsic excitability in LPAG. (A) Representative traces of action potentials (AP) in +/+ and Δ2/+ mouse. (B) AP frequency of LPAG neurons in +/+ and Δ2/+ mouse in response to increasing depolarizing current. (C-E) AP amplitude (C), AP half-width (D), and afterhyperpolarization (E) of the first AP evoked by current injection are comparable between +/+ and Δ2/+ mice. In B-E, the value are presented as mean ± SEM (+/+, n=20/4; Δ2/+, n=33/3). N.S., no significant difference, t-test, SPSS. HW, half-width; AHP, afterhyperpolarization.

**Figure S10.**
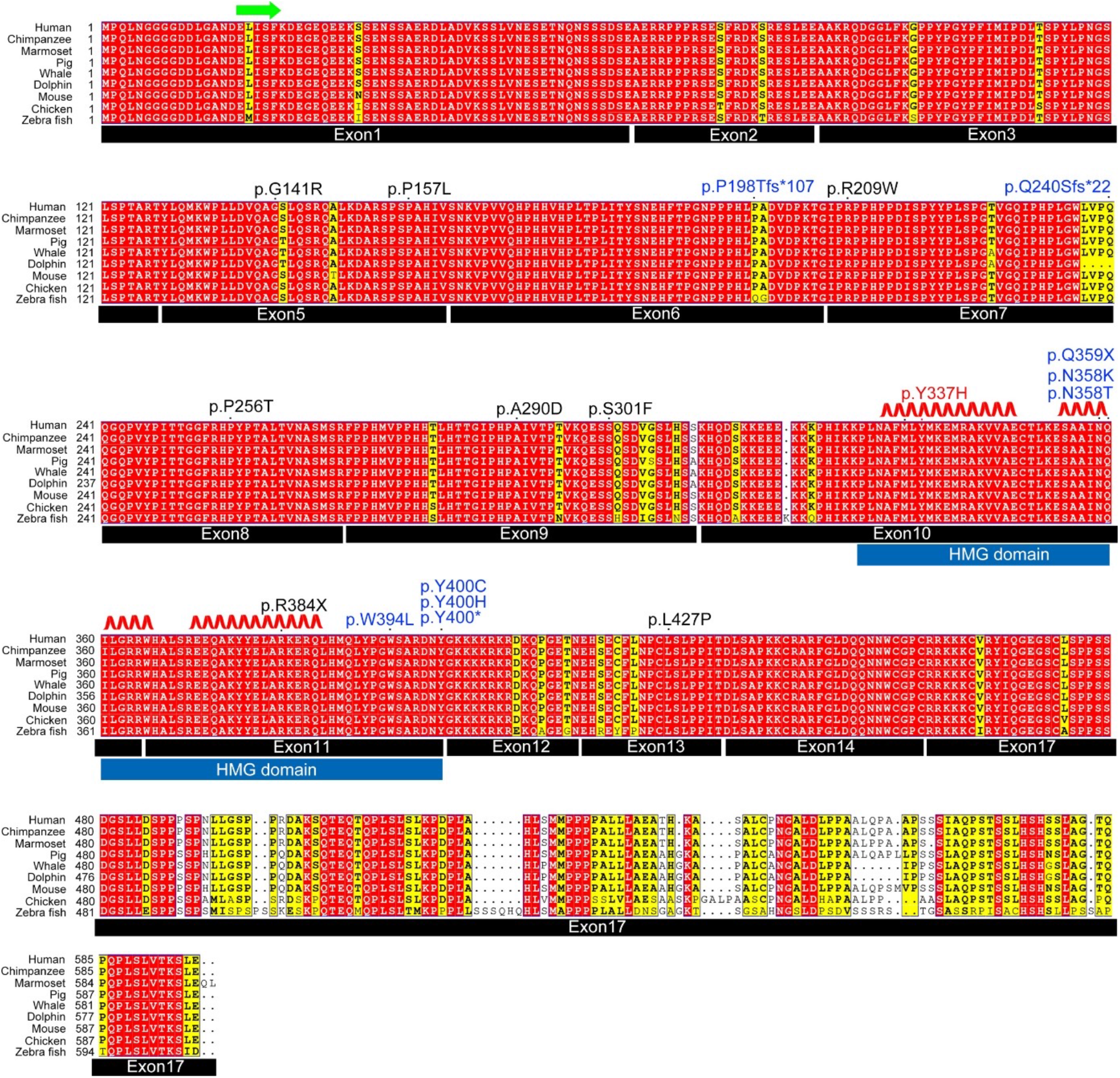
TCF7L2 disease-associated mutations. Protein sequence alignment of TCF7L2 across various species and human TCF7L2 (NM_001198528) were employed for positioning. Disease-associated mutations include: 1) ENU-induced Y337H mutation colored in red; 2) mutations described in previous studies are colored in black (PMID: 25363768, 25533962, and 28191889) and blue (PMID:34003604), respectively.

**Figure S11.**
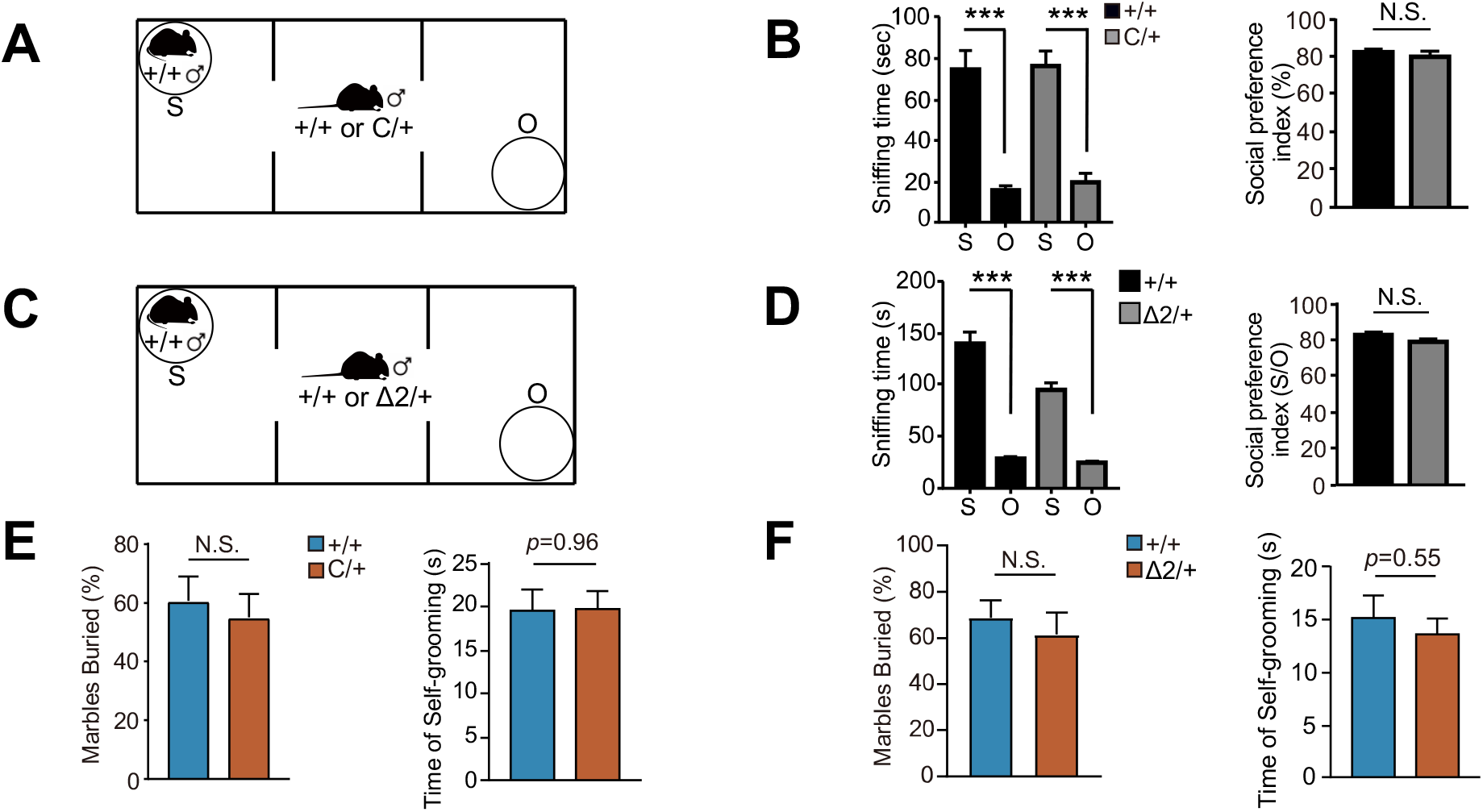
Haploinsufficiency of Tcf7l2 does not lead to ASD-like behaviors in mouse. (A-D) Three-chamber tests of animals with indicated genotypes. Diagrams for the tests (A and C). Social preference index was calculated as previously described (PMID: 31780330). S, stranger; O, object. (E and F) Marble burying test and self-grooming measurement with indicated genotypes. The value are presented as mean ± SEM. N.S., no significant difference, *** *p*<0.001, t-test, SPSS. In B and D, +/+ (n=8-10), Δ2/+ (n= 12), and C/+ (n= 9). In E, for marble burying test, +/+ (n=8) and C/+ (n= 8); for self-grooming test, +/+ (n=16) and C/+ (n= 11). In F, for marble burying test, +/+ (n=8) and Δ2/+ (n= 7); for self-grooming test, +/+ (n=16) and Δ2/+ (n= 14).

**Figure S12.**
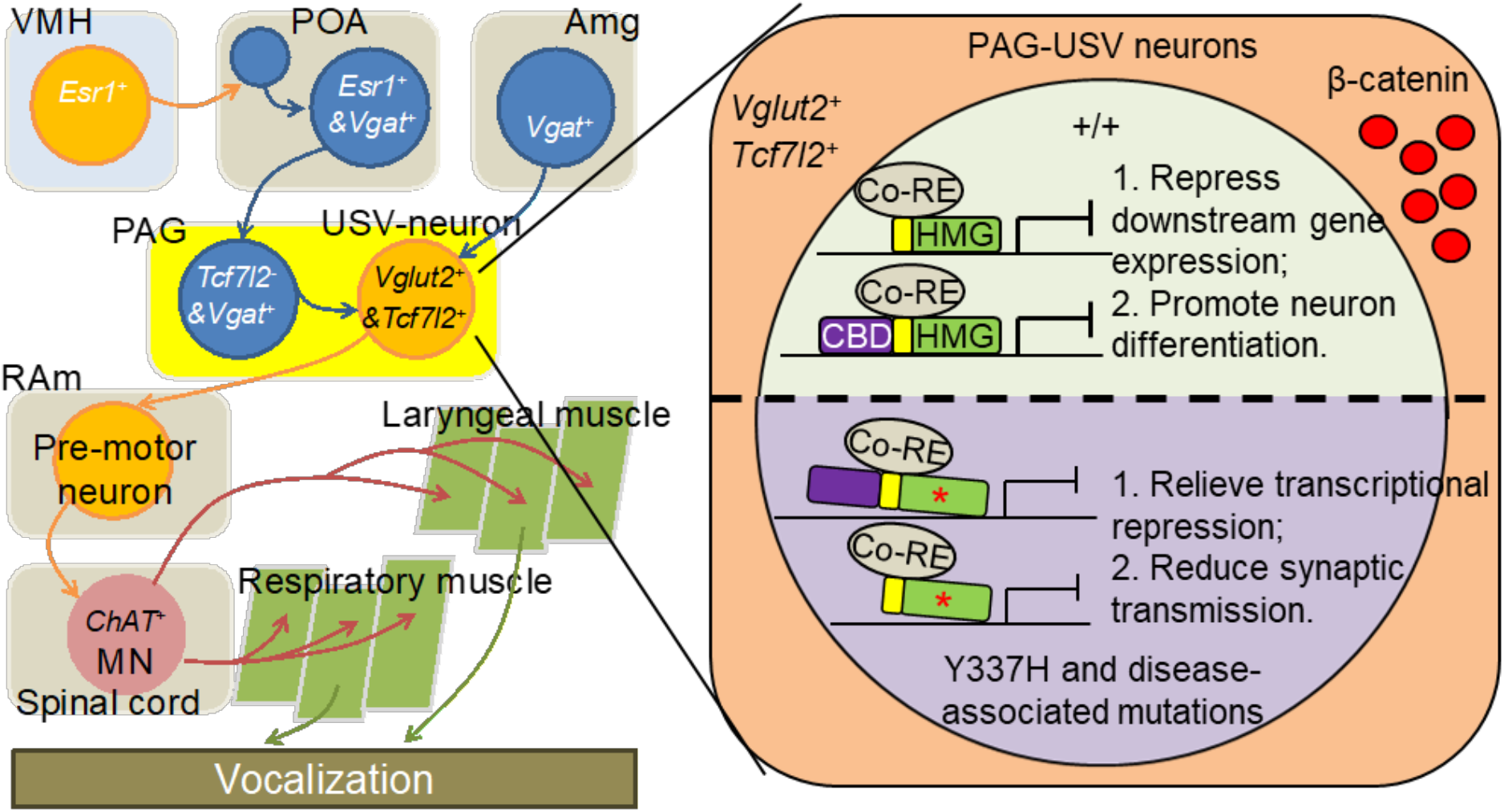
Our working model. Recently, lateral PAG-USV neurons (*Vglut2*+, excitatory) have been identified and characterized, which are necessary and sufficient for mouse USV production (PMID31204083). Besides PAG-USV neurons, *Esr1*-positive neurons in VMH (*Esr1*+) and POA (*Esr1*+ & *Vgat*+) and *Vgat*-positive neurons in Amg (*Vgat*+) and PAG (*Vgat*+) act upstream of PAG-USV neurons to contribute to mouse USV production and persistence (PMID: 33790464, 30900143, 33268894, and 33372655). PAG gates the vocal patterning networks located in caudal brainstem, including RAm, which project to vocal motor neuron (MN, *ChAT*+) pool in spinal cord to control laryngeal and respiratory muscles. The PAG to the brain stem hardware is crucial for both innate and learned vocalization. Here, we demonstrate that *Tcf7l2* is expressed in *Vglut2*+ but not *Vgat*+ (inhibitory) neurons in lateral PAG. Expression of *Tcf7l2* in *Vglut2*+ but not *Esr1*+ or *ChAT*+ neurons is essential for mouse vocalization. Removal of *Tcf7l2* in *Vglut2*+ neurons in midbrain impairs the USV production and lateral PAG synaptic transmission (Left). In PAG-USV neurons, both flTCF7L2 (containing CBD and HMG domains) and dnTCF7L2 (only containing HMG domain) are expressed and required for mouse USV production, suggesting that a transcriptional repression mechanism of TCF7L2 is crucial for mammal vocalization. We speculate that the transcriptional repression promotes neuronal differentiation. However, loss-of-function mutation Y337H and other disease-associated non-synonymous mutations in HMG domain (* in red) relieve the transcriptional repression and reduce neuronal differentiation in these neurons, which in turn decrease synaptic transmission in PAG-USV neurons and leads to impairments of vocal production and syllable complexity (Right). Abbreviations: PAG, periaqueductal gray; VMH, ventromedial hypothalamus; POA, preoptic area; Amg, amygdala; RAm, nucleus retroambiguus; Co-RE, co-repressors.

**Table S1.** Previously reported TCF7L2 mutations associated with ASD, DD, and speech delay by exome sequencing.

| Chr. position | Ref. | Mutation | Transcript | Category | Patients | rs ID | ASD | Speech delay | Source | ExAC | GnomAD | ESP6500 | 1000G |
| --- | --- | --- | --- | --- | --- | --- | --- | --- | --- | --- | --- | --- | --- |
| Chr10:114901076 | G | c.616+1G>A | NM_001198528 | 5' SS | 1 | rs124771269<br>0 | ASD | NA | PMID:25363768 | NA | NA | NA | NA |
| Chr10:114910883 | G | c.932+1G>A | NM_001198528 | 5' SS | 1 | rs147780910<br>2 | ASD | NA | PMID:25363768 | NA | NA | NA | NA |
| Chr10:114912149 | C | c.1150C>T, p.R384X | NM_001198528 | STOP gain | 1 | NA | DD | NA | PMID:25533962 | NA | NA | NA | NA |
| Chr10:114799823 | G | c.421G>A, p.G141R | NM_001198528 | Non-synonymous | 1 | NA | ASD | NA | PMID:28191889 | NA | NA | NA | NA |
| Chr10:114910819 | C | c.869C>A, p.A290D | NM_001198528 | Non-synonymous | 1 | NA | ASD | NA | PMID:28191889 | NA | NA | NA | NA |
| Chr10:114910852 | C | c.902C>T, p.S301F | NM_001198528 | Non-synonymous | 1 | NA | ASD | NA | PMID:28191889 | NA | NA | NA | NA |
| Chr10:114925397 | C | c.1457C>T, p.S486L | NM_030756 | Non-synonymous | 1 | rs773530340 | ASD | NA | PMID:28191889 | NA | 2.45E-05 | NA | NA |
| Chr10:114925435 | C | c.1495C>T, p.R499X | NM_030756 | STOP gain | 1 | rs375657594 | ASD | NA | PMID:28191889 | 5.01E-05 | 8.23E-06 | NA | NA |
| Chr10:114925475 | C | c.1535C>T, p.S512L | NM_030756 | Non-synonymous | 1 | rs770242240 | ASD | NA | PMID:28191889 | NA | 2.85E-05 | NA | NA |
| Chr10:114849205 | A | c.531_532insG, p.Y177fs | NM_001146283 | Frame shift | 1 | rs756433967 | ASD | NA | PMID:28191889 | NA | 2.45E-05 | NA | NA |
| Chr10:113141183 | G | c.484-1G>A | NM_001198528 | 3' SS | 1 | NA | NOT | Yes | PMID:34003604 | NA | NA | NA | NA |
| Chr10:113152440 | T | c.1200T>G, p.Y400X | NM_001198528 | STOP Gain | 1 | NA | ASD | Yes | PMID:34003604 | NA | NA | NA | NA |
| Chr10:113144024 | C | c.718del, p.Q240Sfs*22 | NM_001198528 | Deletion | 1 | NA | ASD | Yes | PMID:34003604 | NA | NA | NA | NA |
| Chr10:113151867 | C | c.1075C>T, p.Q359X | NM_001198528 | STOP Gain | 1 | NA | ASD | Yes | PMID:34003604 | NA | NA | NA | NA |
| Chr10:113141291 | A | c.591dup, p.P198Tfs*107 | NM_001198528 | Duplication | 1 | NA | NOT | Yes | PMID:34003604 | NA | NA | NA | NA |
| Chr10:113146097 | G | c.806+1G>C | NM_001198528 | 5' SS | 1 | NA | NOT | Yes | PMID:34003604 | NA | NA | NA | NA |
| Chr10:113151866 | C | c.1074C>G, p.N358K | NM_001198528 | Non-synonymous | 1 | NA | NOT | Yes | PMID:34003604 | NA | NA | NA | NA |
| Chr10:113151865 | A | c.1073A>C, p.N358T | NM_001198528 | Non-synonymous | 1 | NA | NOT | Yes | PMID:34003604 | NA | NA | NA | NA |
| Chr10:113151873 | G | c.1181G>T, p.W394L | NM_001198528 | Non-synonymous | 1 | NA | NOT | Yes | PMID:34003604 | NA | NA | NA | NA |
| Chr10:113152438 | T | c.1198T>C, p.Y400H | NM_001198528 | Non-synonymous | 1 | NA | NOT | Yes | PMID:34003604 | NA | NA | NA | NA |
| Chr10:113152439 | A | c.1199A>G, p.Y400C | NM_001198528 | Non-synonymous | 1 | NA | ASD | Yes | PMID:34003604 | NA | NA | NA | NA |

Note: The chromosome position is based on human reference sequence (GRCh37, 2009). 5' SS, 5' splicing site. ASD, Autism Spectrum Disorder. DD, Developmental Disorder. ExAC, The Exome Aggregation Consortium; GnomAD, The Genome Aggregation Database; 1000G, The 1000 Genomes Project. 1000G: 3,904 smaples (http://www.internationalgenome.org/data-portal/sample). ASD520+Brain547: Chinese 4,224 ASD and DD samples. GnomAD (exome): 119,116 (https://macarthurlab.org/2018/10/17/gnomad-v2-1/).

